# Transcriptomics and chromatin accessibility in multiple African population samples

**DOI:** 10.1101/2023.11.04.564839

**Authors:** Marianne K DeGorter, Page C Goddard, Emre Karakoc, Soumya Kundu, Stephanie M Yan, Daniel Nachun, Nathan Abell, Matthew Aguirre, Tommy Carstensen, Ziwei Chen, Matthew Durrant, Vikranth R Dwaracherla, Karen Feng, Michael J Gloudemans, Naiomi Hunter, Mohana P S Moorthy, Cristina Pomilla, Kameron B Rodrigues, Courtney J Smith, Kevin S Smith, Rachel A Ungar, Brunilda Balliu, Jacques Fellay, Paul Flicek, Paul J McLaren, Brenna Henn, Rajiv C McCoy, Lauren Sugden, Anshul Kundaje, Manjinder S Sandhu, Deepti Gurdasani, Stephen B Montgomery

## Abstract

Mapping the functional human genome and impact of genetic variants is often limited to European-descendent population samples. To aid in overcoming this limitation, we measured gene expression using RNA sequencing in lymphoblastoid cell lines (LCLs) from 599 individuals from six African populations to identify novel transcripts including those not represented in the hg38 reference genome. We used whole genomes from the 1000 Genomes Project and 164 Maasai individuals to identify 8,881 expression and 6,949 splicing quantitative trait loci (eQTLs/sQTLs), and 2,611 structural variants associated with gene expression (SV-eQTLs). We further profiled chromatin accessibility using ATAC-Seq in a subset of 100 representative individuals, to identity chromatin accessibility quantitative trait loci (caQTLs) and allele-specific chromatin accessibility, and provide predictions for the functional effect of 78.9 million variants on chromatin accessibility. Using this map of eQTLs and caQTLs we fine-mapped GWAS signals for a range of complex diseases. Combined, this work expands global functional genomic data to identify novel transcripts, functional elements and variants, understand population genetic history of molecular quantitative trait loci, and further resolve the genetic basis of multiple human traits and disease.

## Introduction

Advances in genome sequencing are providing the opportunity to more completely represent the human genome for the global population^1,2^. For example, recent analyses reporting 300 Mb of previously undescribed genomic sequence in the reference was identified from primarily African-American and Afro-Caribbean samples^2^. To complement these advances and map functional genomic elements and variant effects in these regions requires corresponding functional maps in diverse populations^3–6^. Currently, these resources have been limited^7–10^. Given the increased genomic diversity and lower linkage disequilibrium^11,12^ in African populations relative to other global populations, functional genomic studies provide increased opportunity to map functional elements, identify functional variants, and understand their roles in complex traits and disease^13–16^.

To address the need for large-scale functional genomics resources from diverse populations, we generated gene expression data using RNA-Seq in lymphoblastoid cell lines (LCLs) from 599 donor individuals across six African populations from the 1000 Genomes^12^ and HapMap^17^ projects. We measured chromatic accessibility using ATAC-Seq on a subset of 100 individuals selected from across the six populations. These data allowed us to quantify novel transcription from unannotated regions of the reference genome, as well as from recently described genomic regions not currently in the reference genome^2^. We identified single nucleotide polymorphisms (SNPs) associated with gene expression (expression quantitative loci, eQTLs), alternative splicing (splicing quantitative loci, sQTLs), and chromatin accessibility (chromatin accessibility QTLs, caQTLs), as well as structural variants (SVs) associated with gene expression (SV-eQTLs). We further characterized patterns of population sharing and selection among the QTL variants. Finally, we developed machine learning models to predict functional consequences of variants on chromatin accessibility. Integrating these RNA-seq and ATAC-seq data further enhances the interpretation and fine-mapping of causal variants from GWAS. This approach significantly contributes to our understanding of genome function by expanding the number of genetic variants with a functional annotation in ENCODE4. Together, these data form an African Functional Genomics Resource (AFGR) that aids in elucidating functional genomic elements and genetic variants involved in complex traits and diseases across global populations.

## Results

### RNA- and ATAC-sequencing in six African population samples

Six African populations were represented in the samples selected for RNA-Seq: 99 Esan in Nigeria (ESN), 112 Gambian in Western Division - Mandinka (GWD), 97 Luhya in Webuye, Kenya (LWK), 83 Mende in Sierra Leone (MSL), and 42 Yoruba from Ibadan, Nigeria (YRI) from the 1000 Genomes Project^12^, and 166 Maasai in Kinyawa, Kenya (MKK) from the HapMap Project^17^ (**Figure 1a, Supplementary Table 1**). RNA libraries were extracted from LCLs and sequenced using 75 bp paired end sequencing to an average library size of 42.2 million reads and mapped to GRCh38 (**Methods**). Whole genome sequencing for ESN, GWD, LWK, MSL and YRI was accessed through the 1000 Genomes Project^12^ and completed for MKK (**Methods**). To assess the relationship between genetic variation and chromatin accessibility, we performed ATAC-Seq in a subset of 100 samples distributed evenly among the six populations (**Methods**).

**Figure 1.**
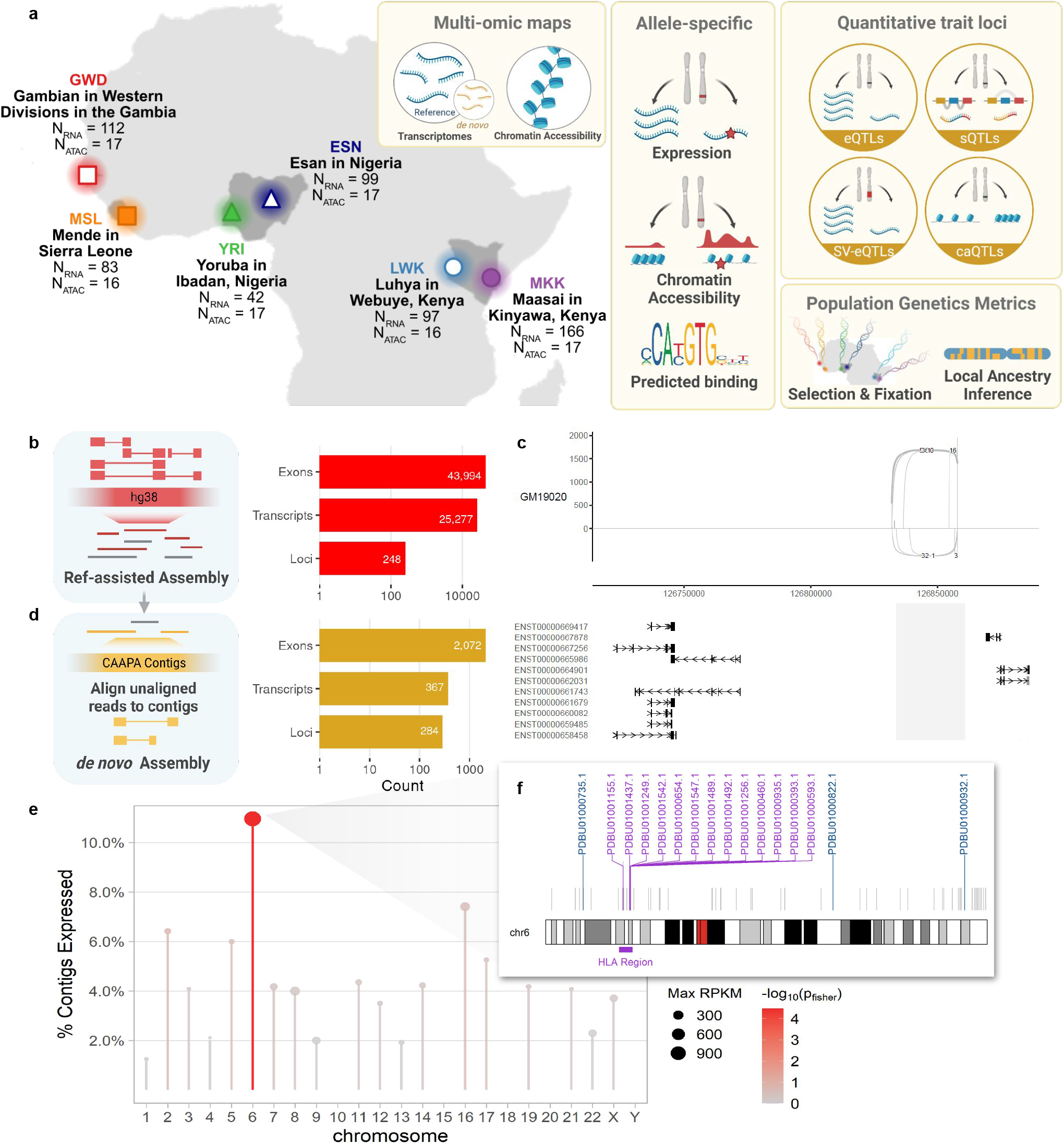
Study overview and transcriptome diversity in African population samples. a. Map shows the number of donors from each population with RNA-Seq and ATAC-Seq samples in this study. Boxes highlight the primary data types made available with this resource: enhanced transcriptome annotations, open chromatin maps, observed allele-specific expression and chromatin accessibility measures at heterozygous sites, predicted allele-specific chromatin accessibility at all sites, quantitative trait loci for expression, splicing, and chromatin accessibility, iHS and Fst scores for the top percent of SNPs under selection, and local Eurasian ancestry estimates for all genes. b. Number of novel exons, transcripts, and loci detected from the reference-aligned reads present in at least 5% of samples, but not in the GENCODEv27 reference or NA12878 long read RNA-seq data^19^. “Loci” refers to distinct transcript clusters. c. Sashimi plot of an example novel multi-exon locus from reference-aligned transcripts mapping to chr12:126831804-126858028 between LINC00944 and LINC02372 (bottom). d. Number of exons, transcripts, and loci detected from reads aligning to the CAAPA contigs from Sherman et al^15^. e. Percent of CAAPA contigs anchored in each chromosome that demonstrated expression in the AFGR RNA-seq data. Lollipop size indicates the maximum expression level across all CAAPA contig-aligned transcripts on the given chromosome. Color indicates -log10 of the Fisher Exact p-value. f. Ideogram showing locations of the CAAPA contigs anchored to chromosome 6. Grey, unlabeled lines indicate the insert positions of unexpressed contigs. Contigs with at least one expressed transcript are colored and labeled with the contig accession number; blue indicates expressed contigs outside the HLA region and purple indicates expressed contigs within the HLA region.

### Novel transcripts identified from African population samples

We identified isoforms in AFGR transcriptomes not present in GENCODE annotations^18^ for GRCh38 using reference-guided transcriptome assembly (see **Methods, Extended Figure 1a**). We detected 57,916 exons, 26,521 high-confidence isoforms of known genes, and 1,303 clusters of transcripts that map to the same locus (potential genes) not previously annotated in GRCh38 GENCODEv27 annotation. We intersected these potential novel features with a newer GENCODE reference (v43), and long-read transcriptome data from NA12878^19^ to determine which features had not been reported in more recent resources, resulting in 24,032 transcripts and 43,400 exons not present in the GENCODEv43 annotation or NA12878 long read transcriptome spanning known genes (**Supplementary Table 2**). Furthermore, we identified 248 novel potential genes, supported by 1245 transcripts and 594 exons (**Figure 1b**). An example of a novel multi-exon locus mapping to chr12:126831804-126858028 between LINC00944 and LINC02372 is shown in **Figure 1c**. Of the 248 potential genes, six overlapped dosage-sensitive regions of the genome implicated in disease^20^. These dosage-sensitive regions tend to be gene-rich, with identifiable dosage-sensitive driver genes; three of the six regions with an AFGR potential gene contain a known haploinsufficient gene, but the remaining three are less well characterized.

### Novel sequences transcribed from genome contigs missing from GRCh38

To mitigate the impact of historical curation biases towards European-derived transcriptomes^7,21,22^, we compiled an enhanced reference transcriptome, incorporating *de novo* assembly of transcripts aligning to 300Mb of contiguous genomic sequences (contigs) identified in individuals of African descent but missing from the current reference (GRCh38)^2^. We detected 284 distinct, transcribed loci specific to these sequences, supported by 2,072 exons across 374 transcripts, totaling 364.9kb of previously uncharacterized transcribed sequence (**Figure 1d, Extended Data Figure 1b**). This transcription from regions not in the current GRCh38 reference genome highlights the need for representative reference genomes for more complete characterisation of diverse transcriptomes and the functional regions of the genome.

The detected transcripts mapped to 266 unique contigs, of which 57 had at least one end anchored in the GRCh38 reference genome (**Extended Data Figure 1b inset**), accounting for 18% (66.76kb) of the transcribed non-reference sequence (**Extended Data Figure 1c**). Although the majority of transcripts fall near either the start or end of their parent contig and thus may reflect partial transcripts (**Extended Data Figure 2c**), we still observe transcripts up to 3kb with a negative correlation between median expression values and transcript size with some exceptions (**Extended Data Figure 2a, 2b**). Contigs that mapped to known genes were enriched for expression over intergenic and unmapped contigs (two-sided Fisher’s exact test; p-value = 2.03×10^-5^, and p-value = 2×10^-16^, respectively **Extended Data Figure 1b**). Chromosome 6 was particularly enriched for contigs with evidence of expression (two-sided Fisher’s exact test; p-value = 2.2×10^-6^), harboring 16 of the 57 contigs (**Figure 1e**). The majority of these chromosome 6 expressed contigs mapped to the HLA region with expression in all populations (13/16, **Figure 1f, Extended Figure 2d**), suggesting uncharacterized genetic and transcriptional diversity in this key immunoregulatory region.

### Consistent patterns of gene expression and chromatin accessibility across populations

Reflecting prior work^23^, we found population structure impacting functional sites explains a limited amount of total variation, as the first two principal components (PCs) for gene expression read counts (**Figure 2a**) and ATAC-seq peak counts (**Figure 2b**) did not separate samples by population. By contrast, the first two genetic PCs differentiated populations by geography and global (genome-wide) Eurasian admixture, respectively (**Figure 2c**). Only a small number of genes or chromatin regions were both significantly different between populations (adjusted p < 0.01) and passed an absolute log-fold-change threshold >1.5 (**Methods**). Out of 11,465 possible genes, we identified just 15 population-specific differential expression events using a linear mixed effects model with nested fixed-effect terms for collection date and population (**Methods, Supplementary Table 3**). Similarly, only three of 95,171 chromatin accessibility regions showed population-specific accessibility patterns under this model (**Supplementary Table 4**).

**Figure 2.**
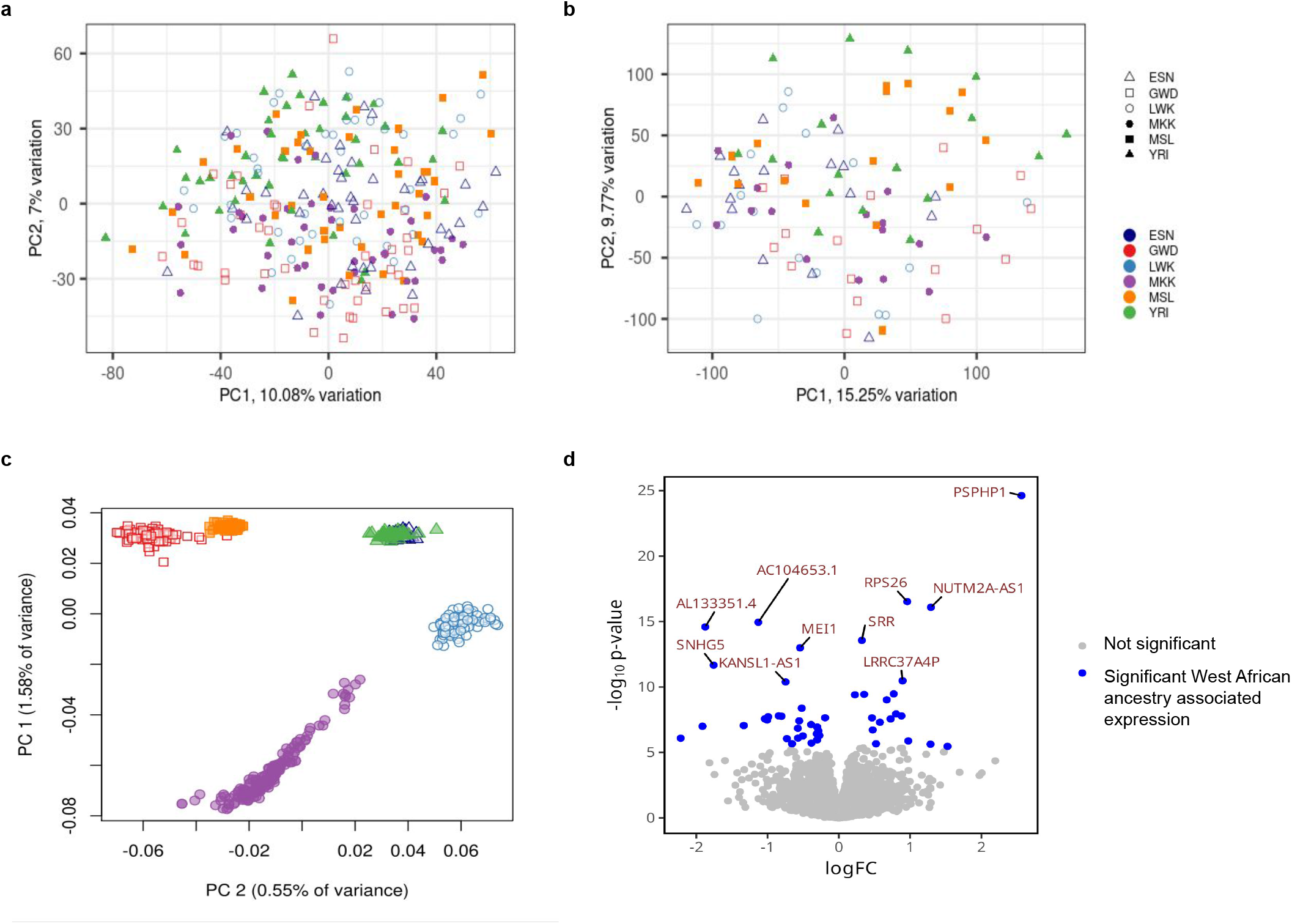
Analysis of gene expression, chromatin accessibility and genetic differentiation by population and local ancestry. a. Principal component analysis (PCA) of gene expression by population. The first two principal components explain 10.08% and 7% of the variation, respectively. b. PCA of chromatin accessibility by population. The first two principal components explain 15.25% and 9.77% of the variation, respectively. c. PCA of genetic differentiation by population. The first two principal components explain 1.58% and 0.55% of the variation, respectively. d. Volcano plot of gene expression vs local Yoruban ancestry inference.

### Global and local ancestry associated with gene expression within Africa

To further investigate ancestry-associated patterns of gene expression, we evaluated the relationship between gene expression and local (within 100 kb of gene) or global (genome-wide) ancestry (see **Methods**). West African haplotypes were defined as those that were more Yoruba-like relative to European (CEU, TSI), East Asian (CHB, JPT), and South Asian (BEB, STU) reference populations from the 1000 Genomes Project (see **Methods**). We identified 567 genes whose expression was correlated and 826 genes whose expression was inversely correlated with global West African ancestry (FDR < 0.05, **Supplementary Table 5**). We also identified 120 genes whose expression was directly correlated with local West African ancestry, and 128 genes inversely correlated with local West African ancestry (FDR < 0.05, **Figure 2d, Supplementary Table 6**).

Expression of the gene with the strongest local ancestry association, PSPHP1 (a.k.a. PSPHL; FDR = 2.22e-20, effect size = 2.53), has been correlated with severity of several diseases including Fanconi Anemia^24^ and sex-hormone derived cancers^25^. A population-stratified 30kb deletion of the PSPHP1 promoter that prevents expression of the gene is rarer in African populations, resulting in higher expression of PSPHP1 in African-ancestry populations relative to other global populations, and potentially biased prognosis for African-descent patients if taken as a stand-alone biomarker^25^. We demonstrate that, even within African populations, local ancestry at this region impacts PSPHP1 expression such that individuals with West African-like haplotypes have higher PSPHP1 expression.

### Expression quantitative trait loci (eQTL) discovery and fine-mapping in African population samples yields more independent signals and smaller credible sets

We measured the association between single nucleotide variant or indel genotypes and gene expression, using a linear regression model to identify eQTLs. After filtering lowly expressed genes, the resulting 15,853 genes were tested in each AFGR population separately; we observed between 1,291 and 3,923 genes with eQTLs (eGenes) at a false discovery rate (FDR) of 5% per population, excluding YRI (**Extended Data Table 1**). As expected, we noted fewer total eGenes in YRI due to reduced power from the smaller sample size (N = 281 eGenes; FDR 5%). A meta-analysis conducted across the six African populations included in AFGR (see Cohort Description) to maximize statistical power tested 11,943,316 unique variants and identified 8,881 eGenes; we refer to these as “African eQTLs” in subsequent analyses.

To identify potentially causal variants from nominally associated eQTL variants (eSNPs, p-value < 1e-5), we fine-mapped these African eQTLs and found an average of 6.02 independent QTL signals per gene, when allowing for up to 10 independent signals using SusieR^26^ (see **Methods**). These credible sets had an average size of 2.1 variants, and 74.1% (37,270/50,280) were singleton sets. We leveraged publicly available RNA-seq from the Genetic European Variation in Disease (GEUVADIS) project to create a complementary, uniformly-processed eQTL callset from European-derived LCLs for fine-mapping comparison between two continental groups (see **Methods**). We refer to the meta-analysis of the GEUVADIS European population eQTL sets as “European eQTLs” in downstream analyses. For genes with fine-mapped eQTL signals from both the African and European eQTLs, the African eQTLs generated more independent signals (**Figure 3a**) and smaller credible sets (**Figure 3b**), with 74.2% (33,651/45,339) of the credible sets containing a single variant, compared with 16.6% (817/4,916) from the European-derived summary stats. Similar results were observed at population-level eQTLs (**Extended Data Figure 3**).

**Figure 3.**
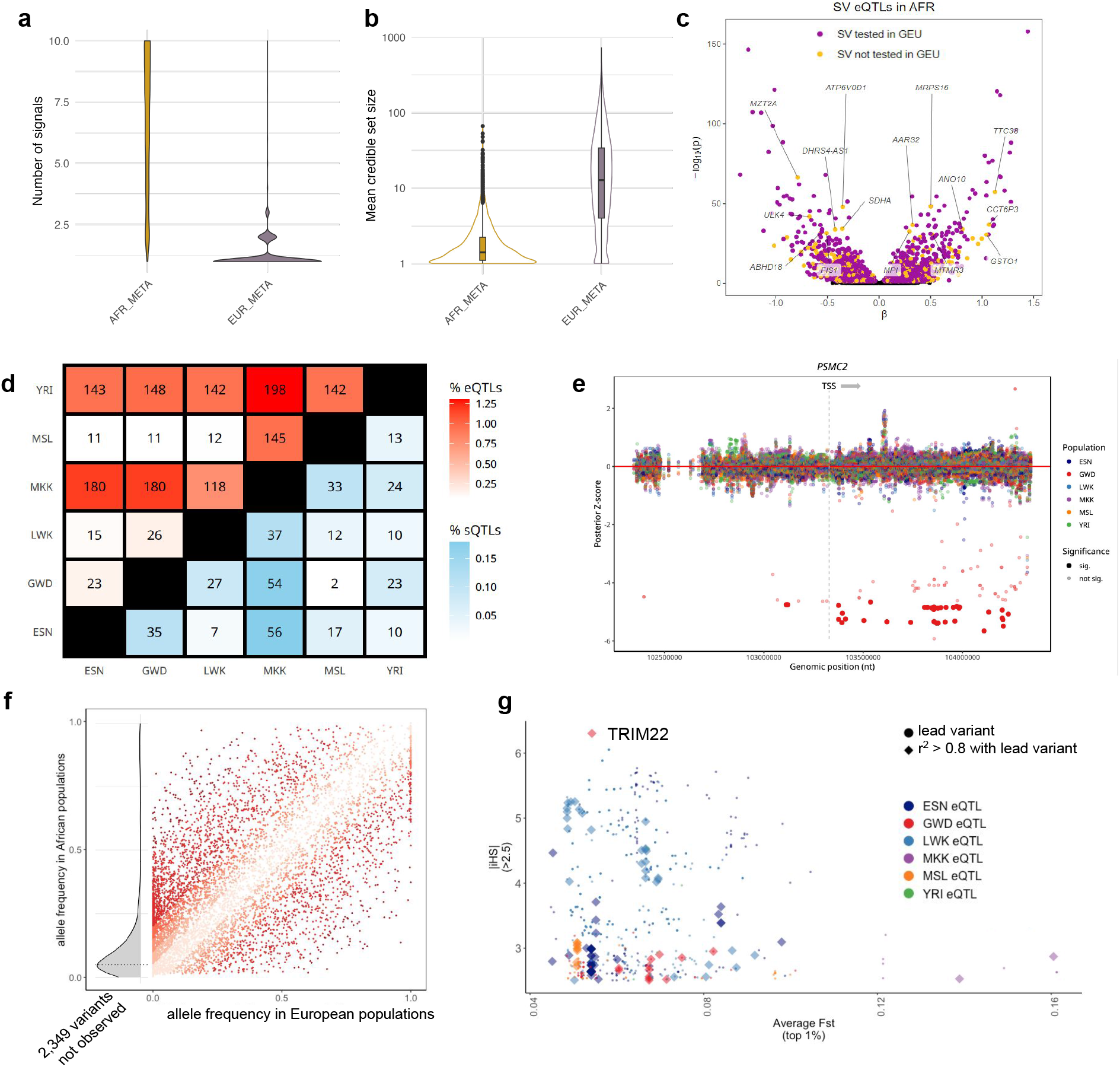
Overview of quantitative trait loci results. a. Number of independently associated signals per gene in the African and European meta-analyzed eQTLs, for genes with a signal in both groups. b. Mean credible set size for African and European meta-analyzed eQTL genes, where genes had a signal in both groups. c. Volcano plot of SV eQTLs and the estimated effect of the alternative allele on expression (β). Significant SV-eQTLs (10% false discovery rate) tested in both AFGR and 1000 Genomes European samples from GEUVADIS (GEU) are colored in purple; significant SV-eQTLs tested only in AFGR are colored in yellow and labeled with the eGene name. d. Number of QTLs with a significant difference in effect size between the indicated populations, determined using deviation contrasts derived from *mashr*. The color gradients represent the proportion of the eQTLs or sQTLs with significantly different effects, as a percentage of the total number that were tested in both populations. e. Example of a population-specific PSMC2 eQTL in GWD, with mashr-derived posterior z-scores representing the effect size of variants in this region of chromosome 7 in each population. f. Allele frequency distribution of lead AFGR meta-analysis eQTL variants in African and European populations from the 1000 Genomes Project. The allele frequency distribution in AFGR of lead meta-analysis variants not present in European populations (n=2,349) is depicted in gray at the far left of the plot. g. Lead variants (diamond), and variants in high ld (r2 > 0.8) with lead variants (circle), from population-level eQTLs with |iHS| > 2.5 and top 1% average Fst scores.

### Structural-Variant eQTLs (SV-eQTLs), splicing QTLs (sQTLs) and chromatin accessibility QTLs (caQTLs) provide additional functional annotation

To measure the impact of structural variants (SVs) on gene expression in AFGR, we tested for associations between gene expression and SVs within 1 Mb from the TSS across all 1000 Genomes populations in AFGR (ESN, GWD, LWK, MSL, YRI) together (see **Methods**). Genotypes were determined from short reads using the graph-based SV genotyper Paragraph,^27^ on a dataset of 107,866 SVs discovered from long-read sequencing of 15 individuals (3 of which possessed predominantly African ancestries)^28^. Among the 39,201 common African SVs (MAF ≥ 0.05) tested, we identified 2611 SV-eQTLs at an FDR of 5%, of which 516 SVs were not present or low-frequency (MAF < 0.05) in the European samples (**Figure 3c**).^28,29^

We next identified splicing QTLs (sQTLs) with a MAF > 0.05 using LeafCutter^30^ and discovered between 473 to 842 sQTLs in each population at an FDR of 5%, excluding YRI due to the reduced sample size (**Extended Data Table 1**). Similar to eQTLs, we conducted a meta-analysis of sQTLs across AFGR and identified 6,949 sQTLs. To determine how frequently splicing events may impact total gene expression, we tested for colocalization of the top eQTL and sQTL signals per feature using Coloc^31^. We found 346 eGenes from our eQTL meta-analysis with strong evidence of eQTL-sQTL colocalization between the top QTL signals, indicating possible shared genetic architecture underlying both the expression and splicing of the genes (posterior probability >= 0.8, see **Methods**).

Using ATAC-seq data, we investigated chromatin accessibility QTLs (caQTLs) to identify genetic effects on transcription factor binding that could subsequently modify gene expression. We assessed the relationship between genetic variation and chromatin accessibility chromatin accessibility QTLs (caQTLs) with a MAF > 0.05, and identified 10,569 caQTL regions across the genome (see **Methods**). We identified 778 meta-analysis eGenes that colocalized strongly with at least one open chromatin region (posterior probability >= 0.8), representing events where common genetic variant regulation of gene expression may be mediated by chromatin accessibility and transcription factor binding.

### Most eQTLs and sQTLs are shared between populations

We found limited evidence of population-dependent QTL effects, with just a small number of feature-variant pairs showing significant difference in effect size between populations when using Bayesian multivariate adaptive shrinkage with *mashr* to infer posterior variance-covariance matrices between groups^32^. Using deviation contrasts, we identified unique effects in one population compared with all others (**Supplementary Figure 1**). Using pairwise population contrasts, we found that just 0.072% to 1.295% of eQTL and 0.006% to 0.178% of sQTL feature-variant pairs identified in more than one population showed a significant difference in effect between groups (**Figure 3d**). These findings indicate a high degree of sharing of common variant effects between populations, or a lack of statistical power to detect smaller differences in effect size. One example of a gene that showed differences in variant effect sizes among tested populations was *PSMC2*, involved in ATP-dependent degradation of ubiquitinated proteins,^33^ which contained several variants significantly associated with gene expression in GWD (**Figure 3e**). In other instances, more than one population showed evidence of non-zero effect sizes, and the magnitude of the effects significantly differed between populations (**Supplementary Figure 2**). Gene set enrichment analysis did not identify biological pathways overrepresented in the set of genes which contained population-dependent differences.

Consistent with our findings in shared QTL effect sizes, we found population-by-population comparisons yielded replication proportions (pi_1_ values) ranging from 0.63 to 0.94 for eQTLs and 0.41 to 0.70 for sQTLs using Storey’s method,^34^ which uses the distribution of p-values to estimate the proportion of eQTLs in one population that are also eQTLs in a second population. (**Supplementary Figure 3**). For this population-by-population comparison, we excluded YRI, as its smaller sample size and subsequently lower number of QTL discoveries confound interpretation of replication, which more likely reflects power differences than biologically relevant differences.

We further assessed the replication of eQTLs discovered in AFGR with those discovered in the GEUVADIS European samples. pi_1_ values ranged from 0.71 (MSL) to 0.92 (YRI) for replication (**Methods**). In this case, the higher replication of eQTLs detected in YRI is likely driven by the detection of high-frequency eQTLs due to its smaller sample size. For most of the populations (ESN, GWD, LWK, MSL, YRI), about 55% of the all the variants tested for eQTLs in AFGR were also tested in GEUVADIS European populations; about 62% of the variants tested in MKK were also tested in GEUVADIS European populations, consistent with more recent Eurasian admixture in MKK (**Supplemental Figure 5**). The proportion of significant SV-eQTL associations (5% FDR) in the African populations that replicated in the GEUVADIS European population samples was 0.66 (bootstrap 95% CI: [0.59, 0.71]), and did not differ between insertions and deletions. This pi_1_ estimate is slightly lower than those for SNPs, likely reflecting increased uncertainties in SV genotyping relative to SNPs^27^.

### Evidence for selective pressure on eQTLs and sQTLs

To identify eQTLs and sQTLs where selective pressures may have resulted in changes in allele frequency between continental populations, we compared the allele frequencies of meta-analyzed eQTLs and sQTLs in AFGR with the corresponding allele in European populations, as reported in the GEUVADIS population samples (**Figure 3f**). Of the AFGR meta-analysis eQTL lead variants, 2,476 are extremely rare (0 - 0.1% frequency) or missing in high coverage genotypes from European populations in the 1000 Genomes project, 635 are rare (0.1 - 1% frequency) and 666 are low frequency (1% - 5%). This likely represents a combination of population demographic history, including population bottlenecks and genetic drift.

Structural variants with high frequencies among African populations may also be enriched for targets of historical positive selection. We observed that 45.8% of highly differentiated SVs^28^ tested in our study were also nominally significant SV-eQTLs at 5% FDR (compared to 25.9% for all SVs) (**Supplementary Table 7**). Some of these variants showed significant associations with multiple genes, such as 1142_NA19240_del, a 4706 bp deletion on chromosome 1, which was associated with the expression of 12 genes, including five linked to disease in OMIM (https://omim.org/) or OpenTargets^35^: *YY1AP1* (MIM: 607860), *MSTO1* (MIM: 617619), UBQLN4, CKS1B, and SYT1.

To better understand selection between populations within the African continent, we assessed iHS, a measure of haplotype homozygosity,36 and Fst, a measure of population differentiation for eQTL variants^37^. We identified 41 eGenes containing lead eQTL variants with high iHS (absolute value > 2) and high Fst (top 1% per population) scores (eQTLs **Figure 3g**, sQTLs **Supplementary Figure 4**, **Supplementary Table 8**). One lead eQTL variant in GWD with a high iHS score, chr11:5698547 (iHS = 6.3), is associated with expression of TRIM22, a known HIV restriction and antiviral protein^38^. In addition, we identified several HLA genes, near which we have also identified expression in the contigs not currently in the reference sequence described above (see Novel sequences transcribed from genome contigs missing from GRCh38). The eQTL gene list is enriched in two immune related GO terms (PantherDB^39^): the interferon-gamma-mediated signaling pathway, as well as antigen processing and presentation of exogenous peptide antigen via MHC class II.

### Allele specific expression (ASE) identifies rare protein-truncating variants that escape nonsense-mediated decay

Allele specific expression (ASE) is a measure of allelic imbalance that can reveal within-individual regulatory effects from rare and common variants in a context that is inherently controlled for genetic background and environmental exposure. The high rates of heterozygous sites present in African populations relative to other global populations^40^ enabled testing of allelic effects for variants that are not present in other datasets. Across each individual in AFGR, we tested a median of 14,548 heterozygous sites for ASE, and detected a median of 2,000 significant ASE sites (**Supplementary Figure 5a, Methods**). Of the 189,148 unique sites with significant ASE in at least one individual (representing 17,104 distinct genes), 57.0% of the variants were homozygous (97,571 variants) or not annotated (10,246 variants) in 30x-coverage whole genome sequencing of European populations from the 1000 Genomes project (**Extended Data Figure 4a**).

Furthermore, rare protein-truncating variants are expected to trigger nonsense-mediated decay (NMD), which presents a distinctive allele-specific expression pattern favoring expression of the non-truncating allele^41^. Variants that escape nonsense-mediated decay can give rise to erroneously truncated proteins with pathogenic potential^42^. We were able to assess allele-specific expression patterns at 333 high-confidence, predicted protein-truncating variants (see **Methods**) with low allele frequency in AFGR (MAF < 0.05). The majority (232) of these PTVs exhibited allelic imbalance towards the reference allele, indicative of nonsense-mediated decay, including 59 with almost complete lack of alternate allele expression (refRatio > 0.95, see **Methods**). Furthermore, 62.7% of ASE-tested truncating variants showed evidence of NMD in predicted gain-of-function genes^42^ (see **Methods**). However, about 32% (105) of all the predicted protein truncating variants tested did not show evidence of ASE, including 20 variants in predicted gain-of-function genes (**Extended Data Figure 4b**). For example, we observed a rare variant that is predicted to cause an early truncation of the KAT6A protein exhibit evidence of NMD escape, a mechanism linked to one form of autosomal dominant KAT6A syndrome (MIM: 616268)^43–45^.

Beyond rare protein truncating events, ASE sites can also identify allelic effects of variants with a lower allele frequency than the cut-off used to identify eQTLs in this study. We identified 104,701 low frequency ASE variants (MAF < 0.05) significant in at least one individual, of which 84.9% were homozygous or absent in the 30x coverage of the 1000 Genomes European population samples (see **Methods**). As these low-frequency ASE sites may be tagging a more common, heterozygous eSNP that is driving the allelic imbalance in expression^46^, we identified variants with evidence of ASE in samples that harbored no heterozygous eQTLs for the given gene (see **Methods**). The majority (97,875) had no accompanying heterozygous eSNPs in any of the significant ASE samples, representing allelic expression events too rare for eQTL testing in this cohort (**Extended Data Figure 4c**).

### Allele-specific chromatin accessibility expands annotation of variant impacts

Allele-dependent binding affinity of transcription factors, pioneer factors, and other DNA-binding proteins may result in allele-specific chromatin patterns. To prioritize these sites, we identified heterozygous sites associated with allele-specific chromatin accessibility (ASC). In each individual, we tested a median of 22,615 sites and discovered a median of 4,840 sites with significant allelic associations with accessibility (Supplementary Figure 5). Of the 299,247 unique and significant ASC sites discovered in any individual with ATAC-Seq data in AFGR, 77,783 variants were low frequency in the AFGR ATAC-seq samples (MAF < 0.05) and thus not included in caQTL analyses, and 36.9% of variants were homozygous (110,408 variants) or unannotated (205 variants) in 30x-coverage whole genome sequencing of European population samples from the 1000 Genomes project.

To identify allele-specific chromatin accessibility sites that may be impacting gene regulation, we subsetted to variants within active B-cell enhancers based on activity-by-contact (ABC) maps47. We found that 6,645 (2%) of our significant ASC variants fall within active B-cell enhancers targeting 4,345 distinct genes, of which 25% (1,097) are OMIM genes.

We leveraged our allele-specific chromatin accessibility data to validate caQTLs detected at the population level, and further identify eQTLs that may be impacting expression via chromatin accessibility and binding patterns. We observed that 93% (8,890/9,620) of nominally significant caQTL variants (p-value <= 1e-5) tested for ASC showed significant allelic effects in at least one individual, and 37% (3,572/9,620) were high-confidence ASC variants, observed consistently in at least 5 individuals (see **Methods**). We further observed that 64% (25,956/40,047) of nominally significant eQTL variants (p-value <= 1e-5) tested for ASC showed a significant allelic effect on chromatin in at least one individual, and 1,129 of these were high confidence ASC variants, implicating differential chromatin accessibility as a mechanism for gene expression changes at these loci.

Next, we applied a convolutional neural network model to extend prediction of ASC effects even where it is not possible to quantify ASC empirically. ChromBPnet predicts chromatin accessibility at a base-pair resolution using genetic sequence as input while accounting for Tn5 transposase binding bias (see **Methods**).^48^ Predicted difference in chromatin accessibility is higher on average for alleles that were observed to be significant, high confidence ASC sites than sites that did not exhibit ASC (two-sided Fisher’s exact p-value < 0.001; **Extended Data Figure 5a**). Predicted chromatin accessibility impact was well correlated with observed allelic fold changes from our ASC findings (Spearman r = 0.696; **Extended Data Figure 5b**), and effect sizes of significant caQTLs (Spearman r = 0.693 **Extended Data Figure 5c,d**). Thus, in addition to empirically observed ASC, we provided predictive chromatin accessibility scores for 78.8 million variants, of which 741,537 are expected to have significant allelic effects (adjusted p-value < 0.01, see **Methods**). To our knowledge, this resource constitutes the largest resource to date of allele-specific chromatin accessibility predictions derived from African population samples.

About 1% (7,374/741,537) of variants predicted to impact chromatin accessibility by chromBPNet fall within enhancers with evidence of activity-by-contact functionality in B-cells^47^. These enhancers are predicted to target 9,467 unique genes, 26% (2,507) of which are OMIM genes, effectively doubling the number of target genes with allele-specific binding annotations in linked enhancers compared to empirical ASC alone.

To further interpret the potential impact of variants on transcription factor binding, we annotated motif sites surrounding 77,663 variants with significant predicted allele effects (permuted logFC p-value < 0.001). We identified binding motifs encompassing these variants using TF-MoDISco^49^ on variant importance scores from DeepLift^50^ and utilized TomTom^51^ to link these variant-dependent binding motifs to transcription factors and DNA-binding proteins (**Extended Data Figure 6a**). By applying a TF-MoDISco q-value cutoff of 0.05, we successfully annotated 19,649 variants for which there was evidence of human TF motif disruption, creation, swapping, or affinity change (**Extended Data Figure 6b**).

### Fine-mapping GWAS using functional genome annotations from African population samples

We integrated AFGR QTLs with large-scale GWAS (sample size > 10,000) to link disease-associated loci to causal genes and variants (**Supplementary Table 9**). We prioritized 11 GWAS traits for which LCLs were suggested as a relevant cell type by stratified LD Score regression. Consistent with previous ENCODE studies^52^, traits that showed showed enrichment of heritability within chromatin accessible regions of LCLs were predominantly immune-related traits, including Crohn’s disease,^53^ inflammatory bowel disease,^53^ lupus,^54^ multiple sclerosis,^55^ primary biliary cirrhosis,^56,57^ rheumatoid arthritis,^58^ and ulcerative colitis^53^ (**Extended Data Figure 7a**). Like most large-scale GWAS to date^59^, these studies were performed in primarily European cohorts, demonstrating, as expected, that African population-derived LCLs are also useful for interpretation of GWAS across global studies.

We identified 232 significant GWAS regions that may be mediated by changes in gene expression driven by nearby eQTLs as candidates for colocalization and causal variant interpretation (**Extended Data Figure 7b, 7c**). We found moderate evidence of colocalization in 42% (89/210) of GWAS regions near a meta-analyzed AFGR eQTL, and 39% (73/187) of the regions near a meta-analyzed European eQTL (colocalization probability>0.5) (**Figure 4a**). Notably, 33 colocalizations with AFGR eQTLs were missed or not tested with the European eQTL colocalizations, while just 17 colocalizations were unique to the European eQTLs. A subset of GWAS regions (56) colocalized with both African and European eQTL(s). Additionally, 150 GWAS regions had an AFGR caQTL within 10kb of the sentinel variant, and 34% (51/150) showed evidence of colocalization. A subset of caQTL-colocalized regions (23) also colocalized with at least one meta-analyzed AFGR eQTL, representing loci where the phenotype association may be driven by transcription factor activity mediating the expression of an eGene. The number of colocalized GWAS regions per QTL trait set and the number of multi-QTL colocalized regions are summarized in **Extended Data Figure 7c**. Collectively, 140 colocalized regions were nominated for statistical fine-mapping across 11 traits.

**Figure 4.**
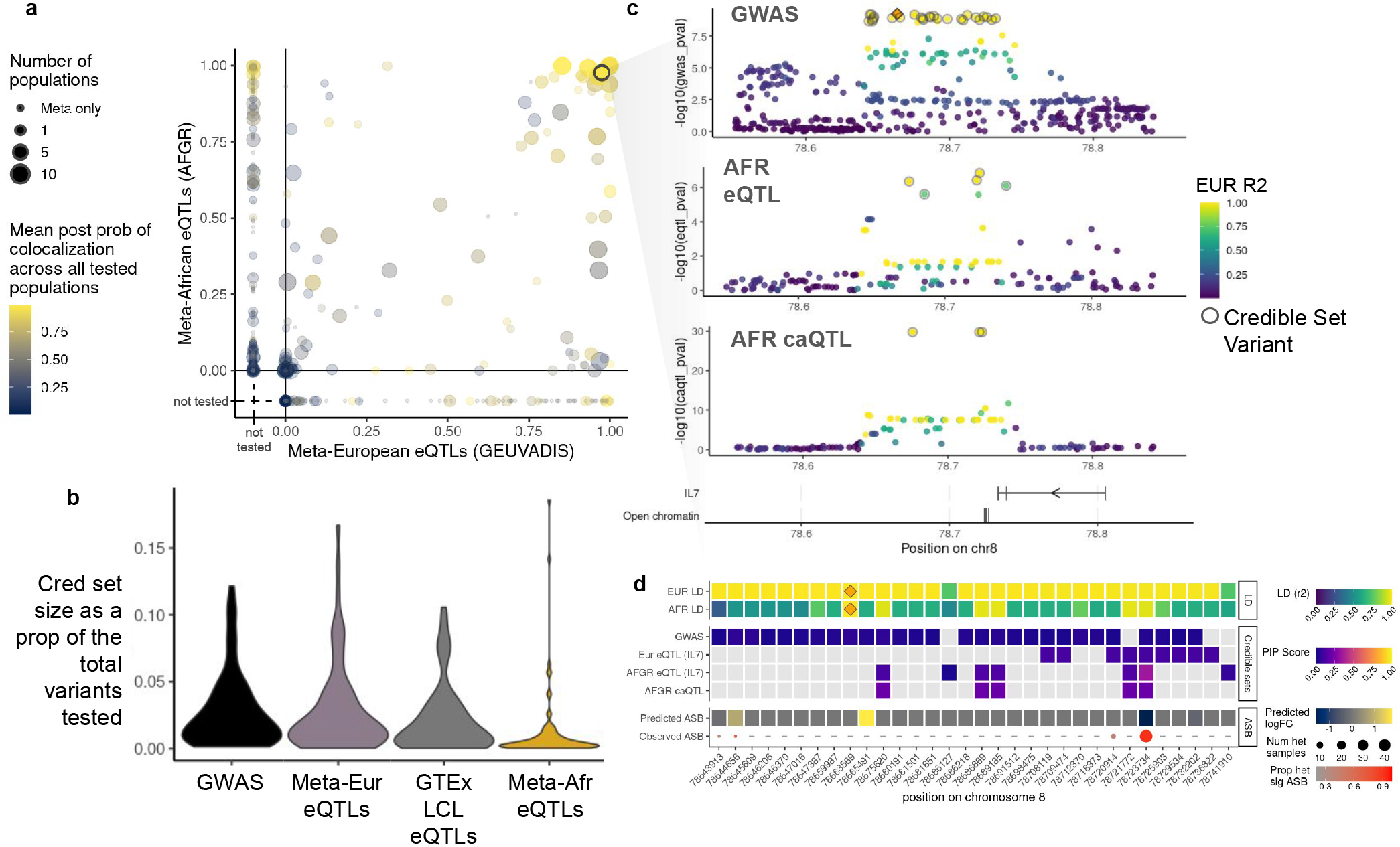
Colocalization and fine-mapping GWAS with QTLs prioritizes functional disease variants. a. Comparison of GWAS colocalizations with the meta-analyzed AFGR eQTLs, and the meta-analyzed European eQTLs from GEUVADIS (y-axis and x-axis, respectively). Loci testable with just AFGR or just GEUVADIS are plotted to the left of x=0 and below y=0, respectively. Point size represents the number of individual eQTL datasets with which the GWAS locus was tested for colocalization. Point color indicates the average probability of colocalization across all the tested groups. The *IL7* locus associated with Multiple Sclerosis is circled in grey. b. Credible set size distributions, presented as a proportion of the total number of variants tested, for GWAS-eQTL colocalized regions. SusieR credible sets were generated for each colocalized region using the GWAS summary statistics and the corresponding colocalized QTL summary statistics independently. c. Locuszoom plots for region on chr8 associated with Multiple Sclerosis (top), expression of *IL7* in the meta AFGR eQTLs (middle), and one of three chromatin accessibility windows near the transcription end site of *IL7* (bottom). Lead GWAS SNP is indicated with the orange diamond in the GWAS panel, and points are colored by LD to the lead variant based on the 1000 Genomes European VCF. Variants prioritized in the SusieR credible sets are circled in grey. The annotation track at the bottom highlights the position and direction of the *IL7* gene, and the genotype-associated open chromatin regions (caQTLs). d. Annotation heatmap of all susieR credible set variants at the *IL7* locus. Top two rows depict African and European LD between credible set variant and lead GWAS variant; middle four rows display the posterior inclusion probability (PIP score) for each variant in the indicated credible set, where light grey indicates that the variant was tested but not included in the credible set; bottom two rows display the predicted allele-specific chromatin accessibility score from chromBPnet, and the proportion of heterozygous samples with significant observed ASC at that variant.

To identify likely causal variants, we fine-mapped each colocalized region using SusieR (140 total regions), assuming a single causal signal for the GWAS and QTL summary statistics independently. The resulting credible sets were subsequently combined to produce a consensus credible of overlapping variants. Overall, we generated GWAS and QTL credible sets for 101 colocalized regions^60^. The median credible set size with the meta-analyzed AFGR eQTLs alone was 2 variants (0.3% of tested eQTL variants per region, **Figure 4b**). We identified a median of 9 credible set variants (1.5% tested variants) for the regions fine-mapped with the meta-analyzed European eQTLs (**Figure 4b**). We observed that, of 51 GWAS loci that colocalized and were fine-mapped with African eQTLs and at least one of the GTEx LCL eQTLs or European eQTLs, 43 loci had smaller credible sets using AFGR than European eQTL sets from GEUVADIS or GTEx.

### African Functional Genomics Resource integration prioritizes functional variant from multiple sclerosis GWAS

To highlight the combined utility of data in AFGR, we explored a multiple sclerosis GWAS region that colocalized with African and European meta-analyzed eQTLs associated with *IL7* expression, an immune-related gene implicated in multiple sclerosis, as well as multiple caQTLs nearby (**Figure 4c**). We examined the credible sets derived from the GWAS and QTL summary statistics; while the GWAS credible set consisted of 28 variants with equal posterior probabilities, one variant, rs2046338, was prioritized across all credible sets, and was in high LD with the sentinel GWAS variant (European R^2^ = 0.99, African R^2^ = 0.95).

We integrated data from across AFGR to further elucidate the potential functionality of this variant, as well as its roles in *IL7* regulation and multiple sclerosis risk. We examined the allele-specific chromatin accessibility logFC predictions generated by chromBPnet, which prioritized this same variant rs2046338, and predicted that the C allele increases chromatin accessibility in the region. We observed significant allele-specific accessibility at this variant in 44 out of 48 heterozygous samples (**Figure 4d**). deepSHAP^61^ scores of the chromBPnet model indicate that the C allele creates a binding motif for OCT family transcription factors (**Extended Data Figure 6**). This variant is located near the transcriptional end site of *IL7* (**Figure 4c**) and publically available Hi-C data indicates active chromatin interaction between the transcription end site and transcription start site of *IL7* (see **Extended Data Figure 8**).^62^ Overall, this implicates open chromatin regions associated with rs2046338 in AFGR as important for *IL7* transcriptional regulation, and demonstrates the utility of this dataset in enhancing prioritization of functional disease variants.

## Discussion

We present an African Functional Genomics Resource (AFGR), consisting of genetic, transcriptomic, and chromatin accessibility data from a collection of African-derived lymphoblastoid cell lines. Leveraging the genetic and functional data of this resource, we expand the goals of ENCODE4 in mapping the functional genome and measuring functional effects of genetic variants. We provide de novo transcript annotations; local and global ancestry inferences across all samples; variant QTL measures for expression, splicing, chromatin accessibility, as well as their iHS and Fst measures; structural variant (SV) genotypes and QTL associations for expression; allele-specific expression and chromatin accessibility annotations, as well as allele-specific chromatin accessibility effects predicted by a machine-learning model.

Our *de novo* transcript annotations expand on the GENCODEv27 reference transcriptome and include expressed gene constructs on an additional 300Mb of genomic sequence derived from African ancestry genomes^2^. We observed an enrichment of uncharacterized expression in the HLA region from the non-hg38-reference sequences, as well as multi-exon and alternatively spliced transcripts mapping to previously unannotated regions of the reference genome, highlighting the importance of studying cell lines from global populations.

Consistent with previous observations that there is more variation within populations than between populations^63^, we do not observe substantial differences in gene expression or chromatin accessibility between the populations included in our study; however, we may be underpowered to detect subtle population differences. Furthermore, though we observed a few QTL signals with differing effects between populations, the majority of eQTLs (>98.7%) and sQTLs (>99.8%) were consistent across groups, indicating that while there are differences in population structure and allele frequency that might impact functional sites, the amount of variation explained at a genome wide gene expression or chromatin accessibility level is small. We did, however, observe an enrichment in high iHS and Fst variants that overlapped eQTL and sQTL associations with immune-related genes, providing supporting evidence for previous observations of immune-related selective pressure perhaps related to differences in infectious disease burden over many generations driven primarily by geographical differences.

By focusing on transcriptomes from genetically diverse populations, we were able to substantially increase variant coverage for QTL testing. We provide nominal QTL statistics for 11.9 million SNPs with a minor allele frequency greater than 0.05 across our 599 samples, including 4.7 million variants that are not present in 30x coverage of European population samples from 1000 Genomes. Furthermore, fine-mapping of eQTLs from African populations yielded more independent signals and more singleton credible sets compared to a similar set of European eQTLs, demonstrating the impact of smaller haplotype blocks and cohort diversity in QTL discovery.

We further assessed allelic-specific expression at 452,865 variants, of which 104,701 were too rare to test for QTLs in AFGR (MAF < 0.05). Additionally, we tested allele-specific chromatin accessibility empirically for 611,603 heterozygous variants, of which 430,465 variants (70.3%) were not assayable in DNaseI hypersensitivity samples from ENCODE4. Finally, we predicted variant effect annotations for 78 million variants, demonstrated high concordance to empirically measured allele-specific chromatin accessibility in our samples, and identified nearly 50,000 variants impacting DNA-binding motifs. Combined, these variant effect annotations provide a large resource of variant annotations can aid interpretation of common and rare disease variants, as demonstrated by the identification of novel and rare protein truncating variants that escaped nonsense-mediated decay.

We further demonstrate the utility of functional genomics derived from African population samples in interpreting GWAS from primarily European populations and exemplify this with a multiple sclerosis locus. By integrating linkage disequilibrium patterns, QTL fine-mapping results, and allele-specific chromatin accessibility information from AFGR, we reduced a GWAS region of 28 equally prioritized credible set variants in perfect LD to a single candidate variant, and identified a potential molecular mechanism by which that variant may impact multiple sclerosis risk through transcription factor binding regulation of *IL7* gene expression. As future GWAS studies expand into non-European populations, we expect resources like AFGR will continue to aid in interpretation of variant effects.

There are limitations to the current resource. The LCLs used here will not fully recapitulate the *in vivo* tissue regulatory environment. However, LCLs remain a useful tool for large scale characterization and this resource expands functional characterization of LCLs beyond pre-existing data from ENCODE and other projects. Crucially, the genetic and functional diversity from these samples represents only a subset of the total diversity across the African continent,^11^ which requires larger and more coordinated efforts to catalog. Our local and global ancestry analyses explore the relationship between gene expression and West African Yoruba-like substructure or broadly Eurasian substructure, but future analyses may benefit from a more nuanced look at different African ancestral components as well. We also note that the GWAS employed in this study are of primarily European descent, in part due to the overwhelming bias towards large European cohorts in generating well-powered genome-wide screens. This bias reflected is reflected in the GWAS easily available for our analyses and results in a failure to evaluate and demonstrate the utility of this resource in mapping the genetic basis of disease in African cohorts or cohorts of recent African descent, where health disparities in genetic and precision medicine are most poignant, and where this resource may have the greatest impact.

Combined, this resource provides new sequencing and variant annotation data from nearly 600 individuals from six communities across Northern Sub Saharan Africa that participated in the 1000 Genomes and HapMap3 projects. This diversity expands our identification of functional elements in the genome, a key goal of the ENCODE project, and enhances the interpretation of both common and rare disease in African populations and globally.

## Methods

### Samples and Genotyping

#### Cohort Description

Individuals from six African populations, including 99 Esan in Nigeria (ESN), 112 Gambian in Western Division - Mandinka (GWD), 97 Luhya in Webuye, Kenya (LWK), 166 Maasai in Kinyawa, Kenya (MKK), 83 Mende in Sierra Leone (MSL) and 42 Yoruba in Ibadan, Nigeria were selected for RNA-Seq. See (**Supplementary Table 1**) for sample IDs, sex, and population assignments.

#### Genome Sequencing and Genotype Calling

We downloaded VCFs aligned to GRCh38 from the 1000 Genomes Project website (http://ftp.1000genomes.ebi.ac.uk/vol1/ftp/data_collections/1000_genomes_project/release/20190312_biallelic_SNV_and_INDEL/, accessed 9 Dec 2020) for ESN, GWD, LWK, MSL, YRI as well as the European populations CEU, FIN, GBR and TSI. Two YRI samples were not present in the vcf provided by 1000 Genomes Project (NA18487 (YRI), NA18498 (YRI)) and excluded from QTL analyses. Four samples missing from the GEUVADIS populations vcf (NA07346 (CEU), NA11993 (CEU), HG00124 (GBR), HG00247 (GBR)) were excluded from downstream analysis, yielding a total of 369 in GEUVADIS samples for eQTL calling. This vcf uses lower coverage sequencing from the original 1000 Genomes release. The accuracy of genotyping in this set is not expected to be an issue for QTL calling with a minor allele frequency > 0.05, as we have computed here. When identifying sites for ASE and ASC, we verified heterozygous sites in vcfs derived from higher coverage (30x) sequencing (see ASE and ASC, below).

A total of 164 Maasai (MKK, HapMap) genomic DNA samples acquired from the Coriell Institute (Camden, NJ, USA) were processed with Illumina PCR-free library preparation and sequenced on an Illumina HiSeq X Ten System in individual lanes (paired end, 30X minimum depth). Two additional MKK samples (NA21486, NA21382), which had been processed with the 10X Genomics Chromium Library preparation, were not included in this batch of WGS and were excluded from downstream analyses that relied on genotype data. Reads were mapped to the GRCh38 reference build and genotype calling was carried out with GATK HaplotypeCaller version 3.8. VCFs from Illumina sequencing were combined with CombineGVCFs, followed by Variant Quality Score Recalibration (VQSR) and phasing with Eagle2 (--outputUnphased flag).

### Structural Variant Genotyping

#### Structural variant eQTL mapping in African and European individuals

To identify structural variant (SV) eQTLs, we intersected gene expression data from this study, and from European individuals in the Geuvadis cohort^3^, with published SV genotypes for the 1KGP individuals^28^. We identified 431 overlapping African samples and 358 overlapping European samples (excluding the YRI population in Geuvadis). Following the SV filtering protocol of Yan et al^28,64^, we removed variants that were rare (MAF ≤ 0.05), violated one-sided Hardy-Weinberg equilibrium (excess of heterozygotes) in more than half of 1K Genomes Project populations, or had a genotyping rate of <50%.

### RNA Expression (RNA-Seq)

#### RNA-Seq Library Preparation, Sequencing and Alignment

RNA was extracted from lymphoblastoid cell lines (LCLs), obtained from the Coriell Institute for Medical Research (Camden, NJ, USA). During RNA preparations care was taken at various steps in order to reduce bias in the final data. The 600 frozen LCLs were processed at Coriell for expansion, followed by pellet preparation, in batches of 100 samples, each containing a balanced representation of all populations. The 600 cell pellets (1e+7) were prepared with RNAprotect (Qiagen) and stored at -80°C until shipment on dry ice. RNA was isolated manually using the RNeasy PLUS mini kit (Qiagen). The cell pellets were assigned to processing batches (12 batches of 48 samples and one of 24 samples) using systematic sampling so that each of the 13 batches for RNA isolation was representative of all populations and of all six pellet batches. The batches of isolated RNA were arranged sequentially in 96-well plates for quality control and preparation of libraries using the standard automated KAPA stranded mRNA library preparation protocol (Roche). One sample was lost in transport before library preparation (HG03478 (MSL)), resulting in a total of 83 MSL samples for RNA sequencing. Stranded RNA libraries (100-300 insert size) were pooled in groups of 12 for sequencing. The libraries in each pool were picked within the same plate through systematic sampling to represent all six pellet batches, the two RNA batches on the plate, and as many populations as possible. The 50 resulting pools were sequenced on duplicate lanes on an Illumina HiSeq 2500 V4 (Paired End 75bp reads). Reads were inspected for quality control using FastQC (http://www.bioinformatics.bbsrc.ac.uk/projects/fastqc), trimmed using Cutadapt^65^ and aligned to GRCh38 using STAR^66^ in two-pass mode, using the GENCODEv27 reference annotation. To check for the possibility of sample swaps, we used verifyBamId^67^; no sample swaps were detected in our experiment (**Supplementary Figure 6, Supplementary Figure 7**).

### Chromatin Accessibility (ATAC-Seq)

#### ATAC-Seq Library Preparation and Sequencing

Cell lines (for list see **Supplementary Table 1**) from 100 donors were received from the Coriell Institute, and cultured 72 hours in RPMI 1640 (ThermoFisher), supplemented with FBS (ThermoFisher), at 5% CO2. Each sample was manually counted using a hemocytometer, and 50,000 cells were aliquoted for ATAC-Seq. ATAC-Seq libraries were prepared according to the protocol of Buenrostro et al^68^ with the addition of Tween (0.1% v/v) to the lysis buffer to reduce the proportion of mitochondrial reads. Libraries were quantified using Qubit (ThermoFisher), as the fluorescence-based system is less sensitive to fragment length distribution than PCR-based methods such as KAPA Quant (Roche). Samples were grouped into pools of 8-10 libraries, and sequenced on an Illumina NextSeq using 75-bp paired-end reads.

#### ATAC-seq Data Processing

The ENCODE DCC ATAC-seq pipeline (http://doi.org/10.5281/zenodo.3662027) (V1.7.0) was used to process bulk ATAC-seq samples, starting from fastq files. The pipeline was executed with “atac.auto_detect_adapter” set to True, while all other arguments were kept at their defaults. The GRCh38 reference genome assembly was used. Biological replicates were analyzed individually, with the FASTQs for the single technical replicate for each biological replicate provided as input to the “atac.fastqs_rep1_R1” and “atac.fastqs_rep1_R2” arguments of the pipeline.

#### ATAC-seq Peak Calling and Quality Control

ATAC-seq peak calling was performed using the most conservative peak set with the smallest irreproducible discovery rate (IDR). In addition, a merged peak set for the entire dataset was generated. The IDR peaks from all biological replicates were concatenated and truncated to within 200 base pairs of the summit (100 bp flanking upstream and downstream of the peak summit). These 200 bp regions were merged with the bedtools^69^ merge command, while keeping the peak summit information from the peak with the lowest MACS2^70^ q-value, yielding 243,588 peaks. The merged peak set was intersected with the IDR optimal peak set for each of the biological replicates using the bedtools intersect command, and only the merged peaks that had at least a 50% overlap in at least 30% of the biological replicates were kept, yielding 71,293 peaks.

To check for possible sample swaps, we used bcftools gtcheck^71^; no sample swaps were detected in our experiment. We excluded GM19468 (LWK), from allele-specific chromatin accessibility and chromatin accessibility QTL analyses, as this sample exhibited a lower insert size distribution and was an outlier in deseq2^72^ normalized counts.

### Detection of Novel Isoforms

To determine whether our Africa Functional Genomics Resource (AFGR) transcriptomes contain previously unannotated transcripts, we employed genome reference-guided transcriptome assembly with StringTie^73^, which uses this mapping to cluster reads, build a model representing all possible isoforms, and estimate the isoforms and their expression simultaneously using a maximum flow algorithm. We then identified isoforms not present in the GENCODEv27 or GENCODEv43 annotation^18^ for GRCh38 using GffCompare^74^.

We used StringTie^73^ to assemble transcriptomes using a genome-guided approach and RNA-seq short reads mapped to the human reference sequence (GRCh38). StringTie We then compared the NA19240 YRI IsoSeq sample^75^ with our YRI samples to determine the coverage that gave the best precision and recall. We found that the optimal coverage was of 0.18 RPKM, which resulted in base resolution sensitivity and precision of 74.5% and 73.8%, respectively, and transcript isoform resolution sensitivity and precision of 49.8% and 59.5%, respectively. We required the transcripts to be observed in at least 5% of the population after filtering for a minimum coverage of 0.18 RPKM. For each sample, we generated a transcriptome annotation in the form of a GTF, and used the StringTie merge method to combine the GTFs into a single transcriptome annotation file, representing a non-redundant, global transcript set across all the samples.

To identify expressed features not present in the annotation used for transcript and gene quantification, we assessed the GTF of global StringTie-constructed transcripts against the GENCODEv27 primary chromosome annotation using GffCompare. We subsequently compared the GTF composed of these 1,303 unannoatated genes with PacBio long-read RNA sequencing from NA12878^19^ with GffCompare and identified those transcripts that had not been previously detected in another assembled transcriptome. Putatively novel features were those that were identified as “unknown/intergenic” by GffCompare in both analyses.

Finally, to determine how many of these features are captured in the most recent GENCODE annotation at the time of publication, we used GffCompare to intersect the set of putatively novel exons, transcripts, and genes (those not present in the GENCODEv27 annotation or the NA12878 long-read RNA-seq data) with the GENCODEv43 annotation for the primary chromosomes (GRCh38). A novel putative gene locus was defined as a distinct cluster of transcripts with shared exons. These novel features were intersected with the dosage-sensitive regions provided from Collins *et al* 2022^20^ using bedtools^69^ intersect.

### Alignment to genetic sequence absent from GRCh38

We aligned all reads to the CAAPA contigs from Sherman et al.^2^ using STAR^66^ two-pass mode, then used Scallop^76^ to create a complete *de novo* transcriptome assembly based on the subset of reads that mapped to the CAAPA contigs. Scallop employs a splice-graph based algorithm to decompose the splice graphs into a minimum number of paths which represent a putative transcript.

Using the de novo assembly, we generated a transcriptome annotation that combines the reference transcriptome and the assembled pan-genome transcripts, and used STAR to realign all reads to the combined transcriptome GTF. Expression was quantified using HTSeq^77^. Putative transcripts from the CAAPA contig *de novo* assembly for which reads mapped more closely to the reference after merging annotations were filtered out of the combined GTF to produce a final enhanced reference transcriptome.

A subset (1,548/125,715) of the CAAPA contigs have at least one end anchored in the GRCh38 reference genome^2^, allowing us to assess where in the current reference genome some of this extraneous transcriptomic information originates from. We leveraged the positional information provided by Sherman *et al* in supplementary tables 1 and 3 to examine trends around which contigs demonstrated expression in AFGR; contigs were categorized according to expression status (“expressed” was defined as at least 0.18 RPKM across at least 5% individuals), and genomic anchor region status (i.e. unmapped, or mapped to an annotated mRNA, exonic, lncRNA, or intergenic region) as inferred from supplementary table 4 from Sherman *et al* (gene intersections for anchored contigs) (**Supplementary Table 2**).

### Principal Component Analysis (PCA)

Principal component analysis was performed separately on the uncorrected RNA-seq, ATAC-seq, and genetic sequencing data for all samples using prcomp in R^78^.

The top ten RNA-seq PCs were tested for correlation with known technical covariates to understand the primary drivers of variation (**Supplementary Figure 3a**). The largest sources of variation significantly correlated with standard covariates. We also noted that PC1 correlated with sample population as a categorical variable, and specifically with the YRI population label, which is the group with the smallest sample size. PC2 correlated with additional population labels, though these populations covary with correlates such as sample collection year. Correlation testing and visualization were performed using ggcorrplot (https://cran.r-project.org/web/packages/ggcorrplot/readme/README.html).

### Differential Gene Expression

To identify genes and chromatin regions that are differentially expressed or accessible by population, we used limma^79^ and DREAM^80^, and filtered out genes in known hypervariable regions.

To compare between populations we subsampled the RNA-seq data such that there were equal numbers of samples per population. Using edgeR, counts were TMM normalized using the function calcNormFactors, converted in CPM values, and filtered such that only genes that passed a CPM threshold of 5 were kept. The non-normalized counts were then transformed and weights were estimated using voomWithDreamWeights. The model was fitted using the function dream.

The mixed model included the following covariates: a random effect for extraction batch and fixed effects for sequencing pool, sex, RNA amount in μg, and RIN. These covariates were identified via a canonical correlation analysis (CCA) as they were not correlated to any other metadata or to the population. We identified a strong batch effect when using this model based on the collection year of each population, with LWK, MKK and YRI having been collected in 2003-2005 and ESN, GWD and MSL having been collected in 2011-2012. Given this substructure, a nested fixed-effects model was used for collection date and population. In this model, a single categorical variable for whether a sample is in the early (2003-2005) or late (2011-2012) batch was used in addition to population specific coefficients for the two populations in each batch other than the referent (early: YRI referent, LWK and MKK alternate; late: ESN referent, GWD and MSL alternate).

We further identified a significant collinearity between populations and culture time. Culture time was computed in units of days by subtracting the recovery date from the harvest date, and ranged between 2 and 25 days. The largest mean difference in culture time between populations was 1.8 days between GWD and MKK, with a mean difference of about 1.8 days, a 53% increase in culture time for the MKK population. We used subsampling to balance mean culture time across populations, choosing up to 11 individuals per population with three days of culture time up to 32 samples individuals per population with four days of culture time, followed by a final subsampling to 40 individuals per population. After accounting for sample collection date as a proxy for Corielle cell line age, time in culture, extraction batch, sequencing pool, sex, RNA amount in μg, and RIN, we identified 15 genes differentially expressed by population with an absolute logFC >= 1.5 and an adjusted p-value < 0.01 (3 ESN, 5 GWD, 1 LWK, 0 MKK, 4 MSL, 2 YRI).

Most of the technical variables in the model are captured within the first ten molecular PCs (**Supplementary Figure 8**). The number of differentially expressed genes detected at a range of logFC and adjusted p-value cut-offs are found in **Supplementary Figure 9a, 9b**.

### Differential Chromatin Accessibility

Population differences in chromatin accessibility was assessed using the same limma DREAM framework as the differential expression. Covariates for the mixed-effects model included population, sex, library concentration, and bead cleaning date as fixed effects and primer 1, primer 2, and sequencing pool as random effects and the same nested fixed effects for collection date and population were used. The number of open chromatin regions detected at a range of logFC and adjusted p-value cut-offs are found in **Supplementary Figure 9c, 9d**.

### Global and Local Yoruba Ancestry

We inferred global ancestry in all individuals using LD pruned (R^2^ cutoff 0.2) variants across all chromosomes with ADMIXTURE^81^ using 7 reference populations including one African population (Yoruba in Ibadan, Nigeria, YRI); two South Asian groups (Bengali from Bangladesh [BEB] and Sri Lankan Tamil in the UK [STU]); two East Asian groups (Han Chinese in Beijing, China [CHB] and Japanese in Tokyo, Japan [JPT]); and two European ancestry groups (Toscani in Italia [TSI] and Utah residents (CEPH) with Northern and Western European ancestry [CEU]). We ran ADMIXTURE with between 2 and 8 clusters and combined the output of these results with CLUMPP^82^. Proportion of global (genome-wide) non-YRI (West African) ancestry for each population is shown in **Supplementary Figure 10**.

We inferred local ancestry estimates in all individuals in 2,295,732 SNPs tested for eQTLs or sQTLs in any population using RFMix with default settings on the phased genomes. We used the same reference populations as those used for global ancestry inference, and a published recombination map^83^. We averaged local ancestry in a 100kb window around each gene to obtain a gene-level estimate of local Yoruba ancestry for each gene in each individual.

Gene expression counts were normalized using the same approach as in the differential gene expression (log normalization of TMM-normalized CPMs). To account for hidden covariates, principal components analysis was run on the counts matrix after normalizing with *corral* (https://bioconductor.org/packages/corral/) followed by feature wise inverse normal transformation. The number of principal components was chosen using the Gavish-Donoho method^84^. The effect of local ancestry on expression was estimated for each gene using a linear mixed effects model with the same covariates as used in differential expression (random effect for randomized extraction batch and fixed effects for sequencing pool, sex, RNA amount in μg, and RIN) In addition we included fixed effects for expression principal components which were not correlated with inverse normal transformed local ancestry at Bonferroni adjusted p < 0.05. The same model was also used for testing association with global ancestry.

We used a linear model to test for differences in local ancestry between populations for each gene. The gene level local ancestry averages used above were normalized with the inverse normal transformation and used as the outcome with population as the predictor. Post-hoc testing of the fitted linear model with *emmeans* used deviation contrasts (one population vs. others) and pairwise contrasts to compare populations.

### Expression Quantitative Trait Loci (eQTLs) Mapping

#### Quantification, Normalization and Hidden Factor Correction

Genes were quantified using HTSeq^77^ and the GENCODEv27 annotation. Lowly expressed genes (genes with a mean of less than five reads across individuals in any population) were filtered, and gene counts were variance-stabilized with DESeq2^72^, then normalized with mean 0 and standard deviation of 1. Multi-allelic variants encoded on one line of the vcf were not able to be interpreted by FastQTL, and dropped from the analysis. We used SVA^85^ to identify hidden factors, and called eQTLs on one chromosome using a range of different numbers of hidden factors to determine maximum sensitivity; the largest number of eQTLs was detected using the maximum number of surrogate variables. In our final eQTLs for each population, we used the maximum number of surrogate variables and genotype PCs explaining cumulative variance up to 5% of the genetic variation as covariates for eQTL discovery. PCs were calculated using SNPRelate^86^. (**Supplementary Figure 11-16. RNA-Seq surrogate variables associated with known variables)**

#### eQTL discovery in African populations

After filtering out lowly expressed genes there were 15,853 genes for eQTL discovery in African populations. Nominal eQTL signals with MAF > 0.05 within 1Mb of the TSS were mapped with FastQTL^87^ on each population separately. We used FastQTL permutation testing, with 1000 to 10000 adaptive permutations with random number seed 123456789 to identify a top eQTL signal for each gene, then implemented Benjamini-Hochberg^88^ procedure to identify genes harboring at least one significant eQTL after multiple testing correction (eGenes). Four samples for which we have RNA-Seq were excluded from QTL analyses as the genotypes were either not present in the VCF from 1000 Genomes (NA18487 (YRI), NA18498 (YRI)) or not sequenced as part of this study (GM21486 (MKK), GM21382 (MKK)).

#### SV-eQTL discovery in African populations

We performed structural variant eQTL (SV-eQTL) mapping with a version of fastQTL modified for use with SVs^87^ (https://github.com/hall-lab/fastqtl), which accounts for SV size when determining the appropriate cis window. As with eQTL mapping performed on SNPs, we conducted nominal and permutation passes, with surrogate variables and genotype principal components accounting for up to 10% of variation between samples as covariates. We used beta-distribution approximated permutation p-values from fastQTL to estimate the false discovery rate (FDR) with the qvalue package^34^ (http://github.com/jdstorey/qvalue). We intersected these significant SV eQTLs with a set of SVs that show substantial allele frequencies differentiation in African populations relative to other global populations, generated by Yan et al.^28^

#### eQTL discovery in GEUVADIS populations

We used RNA-seq data from the GEUVADIS^3^ cohort of samples from across European populations, including Utah Residents (CEPH) with Northern and Western European Ancestry (CEU), British in England and Scotland (GBR), Finnish in Finland (FIN) and Toscani in Italia (TSI). The GEUVADIS study also included 89 YRI samples, of which 21 were also sequenced in our study. Four European population samples, NA07346 (CEU), NA11993 (CEU), HG00124 (GBR) and HG00247 (GBR) were missing from the VCF provided by 1000 Genomes Project and excluded from downstream QTL analyses.

We mapped RNA-seq reads using the same pipelines as for the RNA-Seq from African population samples described above. A total of 15,195 genes passed our expression threshold for eQTL discovery in the European RNA-seq samples from GEUVADIS. Similar to the procedure for African populations, nominal eQTL signals with MAF > 0.05 within 1Mb of the TSS were mapped with FastQTL^87^ on each population separately. We used FastQTL permutation testing, with 1000 to 10000 adaptive permutations with random number seed 123456789 to identify a top eQTL signal for each gene, then implemented Benjamini-Hochberg^88^ procedure to identify eGenes.

### Splicing Quantitative Trait Loci (sQTLs) Mapping

Intron junctions were defined using LeafCutter^30^, for each individual’s STAR two-pass output, and intron excision clusters were determined across all African samples. We filtered out clusters with more than 10 introns. We included introns that were supported by at least 10% of the total number of reads assigned to the cluster in at least 25% of samples in each population and filtered out clusters with fewer than two active introns. To reduce the set to introns with variable usage we filtered out clusters with Hellinger’s distance < 0.01. In total, the filtering steps yielded 13,359 splicing clusters composed of 32,493 intron exclusion events for sQTL discovery. See **Supplementary Table 10** for the number of clusters filtered at each step.

We performed SVA on the filtered intron excision ratios and, as with the eQTLs, tested a range of SVs as covariates with FastQTL to identify sQTLs in the quantile-normalized filtered clusters, for each population separately. For each population, the maximum number of sQTLs was obtained using the maximum number of SVs (4-6), with the exception of YRI, which had the most sQTLs with 1 SV; the maximum number of sQTLs per population were retained for subsequent analysis. The SVs and genotype PCs explaining up to 5% of the cumulative genetic variance were included as covariates in FastQTL. Only biallelic variants were considered as potential sQTLs. We used FastQTL with permutation testing, with MAF > 0.05 and within a 100kb window around each cluster start site, performing 1000 to 10000 adaptive permutations with a random number seed of 123456789. We used Bonferroni correction to adjust for multiple testing across the number of introns in each cluster, followed by Benjamini-Hochberg adjustment for multiple testing over all introns to identify sQTLs with an FDR < 0.05. Finally, we assigned the introns to genes using GENCODEv27.

### Meta-analysis of QTLs

To generate a summary eQTL call set, within-population eQTL nominal p-value summary statistics were meta-analyzed across population groups with the R package mvmeta^89^. All six populations within AFGR were used for the African population meta-analyses: ESN, GWD, LWK, MKK, MSL, YRI. Four populations from GEUVADIS were used for the European population meta-analyses: CEU, FIN, GBR, TSI. We used mvmeta to fit a fixed-effects meta-analysis and produce heterogeneity estimates (Cochran’s Q) for each variant. Only variants at MAF > 0.05 in a population were tested for eQTLs; populations without eQTL summary statistics for a variant were treated as missing in the meta-analysis. Summary statistics for variants with eQTL calls in only one population were carried over directly into the final call set, and were not meta-analyzed. To count the number of significant eGenes in this analysis, we applied eigenMT^90^ to the final eQTL call set, using patterns of linkage disequilibrium as estimated from our samples to estimate the number of effective statistical tests conducted for each gene and adjust for multiple hypothesis testing. The same approach was used to compute a cross-population summary sQTL call set.

We tested 11.9 million variants in the AFGR meta-analysis; for the number of variants tested for eQTLs in multiple large RNA-Seq cohorts, see **Supplementary Figure S17**. Relative to the population-level eQTLs the eQTL meta-analyses identified top eSNPs that tended to be closer to the associated eGene’s transcription start site (**Supplementary Figure S18a,b**), and were better powered to detect significant associations with smaller effect sizes (**Supplementary Figure S18c,d**).

### Replication of GEUVADIS eQTL Discovery

To assess replication of the original GEUVADIS eQTLs results in our re-called datasets, we downloaded the significant GEUVADIS eGene list (ftp://ftp.ebi.ac.uk/pub/databases/microarray/data/experiment/GEUV/E-GEUV-1/analysis_results/EUR373.gene.cis.FDR5.best.rs137.txt.gz, downloaded 11 January 2023), and calculated the proportion which were also eGenes in our bulk-called European eQTL dataset (EUR_GEU), and our meta-analyzed European eQTL dataset (EUR_META). We identified a total of 6,766 eGenes in our meta-analysis, replicating 74.6% of the eGenes in the original GEUVADIS publication (2,432/3,259 eGenes), while identifying an additional 4,334 eGenes. The bulk-called European eQTLs yielded a total of 7,194 eGenes, including 2,506 which were discovered in the original GEUVADIS study, and 5,997 which were also identified in this meta-analysis.

Furthermore, of the 150 eGenes we detected from the 40 YRI samples sequenced for this study after permutation and multiple testing correction, 109 (73%) are also detected in our re-called eQTLs for the GEUVADIS YRI eQTL samples, and 63 (42%) were also detected from the YRI samples in the original GEUVADIS publication. Our re-called YRI eQTLs from the GEUVADIS data yielded 1,510 distinct eGenes after multiple testing correction, in contrast with the 501 detected in the original GEUVADIS analysis; 314 eGenes were reported by both eQTL sets.

### Detection of independent QTL signals

eQTLs were fine-mapped using SuSiE with summary statistics (SuSiE-RSS). The eQTL statistics for all population-level and meta-analyzed AFGR and GEUVADIS datasets were combined with an in-sample LD matrix. For each gene in each data set, all variants tested for nominal association were selected, and pairwise LD (r) was calculated for each pair of SNPs using R’s **cor** function. Any multi-allelic sites or variants missing genotype calls were dropped from the fine-mapping analysis (15% (1742092/11943294) dropped from AFR_META and 0.4% (30824/6965771) dropped from EUR_META) before calculating LD. In addition, variants for which the population was homozygous were dropped. Genes for which SuSiE failed to converge or identified no credible sets were excluded from downstream analysis.

The command for SuSiE-RSS was **susie_rss(bhat = <EQTL_BETA>, shat = <EQTL_SE>, n = num_samples = <NUM_SAMPLES>, R = cor(<FILTERED_GENOTYPES<), estimate_residual_variance = FALSE, check_prior = FALSE, max_iter = 1000, estimate_prior_method = “EM”)**. Default variables set the maximum number of non-zero effects to 10.

We used R version 4.2.2 with susieR version 0.12.35.

### Chromatin Accessibility Quantitative Trait Loci (caQTLs) mapping

The merged IDR peak set (n=243,589) was used for caQTL mapping. We implement SVA on the quantile-normalized read counts, including population covariates. We included all 5 SVs detected, along with the first 5 genetic PCs for caQTL testing using FastQTL. Variants with a MAF > 0.05 in the samples included (subset of 100 individuals) within 10kb of the open region were tested for caQTL using FastQTL with permutation testing, with 1000 to 10000 adaptive permutations and a random number seed of 123456789. The beta-distribution approximated permutation p-values were adjusted for multiple hypothesis testing using Benjamini-Hochberg correction. We also generated nominal p-values of association for all variants using Fastql in nominal mode.

### Population QTL Sharing and Selection

#### mashR

We used mashr^32^ to assess population differences in eQTL and sQTL effect sizes. We examined cases where the feature-variant pair was tested in at least two populations within AFGR, leading to the inclusion of 9,807,451 variants (82.1% of total variants tested for QTLs). Following the strategy in Urbut et al^32^, we extracted a set of strong feature-SNP pairs, and a set of random feature-SNP pairs, because the size of the full dataset was too large to fit the mashr model. The strong feature-SNP pairs were defined as the most significant SNP in each feature in each population (88,586 gene-SNP pairs in eQTLs, 184,248 splice cluster-SNP pairs in sQTLs). Random sets were generated from 1,000,000 random feature-SNP pairs chosen from the eQTLs and sQTLs. The null correlation was estimated from the random set, while the data-driven covariance matrices were estimated from the strong set. We estimated data driven covariance matrices using two methods - extreme deconvolution^91^ from principal components analysis, and flashr.^92^ We used 6 principal components (PCs), the maximum possible number, as input to extreme deconvolution, resulting in 6 covariance matrices as well as a covariance matrix using all 6 PCs for both eQTLs and sQTLs. With *flashr*, we used the default settings for convergence and backfitting which identified 7 factors in eQTLs and 4 factors in sQTLs. We used the recommended approach for choosing factor-specific covariance matrices with non-unique effects by removing any covariance matrix whose associated factor had less than two populations with an absolute loading weight of at least the reciprocal of the square root of the number of populations (1/√6). In addition to the covariance matrix derived from all factors, we retained 2 factor-specific covariance matrices in eQTLs and 1 in sQTLs. The data-driven covariance matrices (10 for eQTLs, 9 for sQTLs) in were combined with 11 canonical covariance matrices and multiplied by a grid of default values of the scaling parameter *omega* (25 for eQTLs, 23 for sQTLs), giving us a final set of covariance matrices (525 in eQTLs, 460 in sQTLs) used to fit the covariance matrix mixture model. The mixture model parameters were optimized on the random feature-SNP set using the default *mixSQP* method.

The fitted mashr models for eQTLs and sQTLs were used to estimate posterior mean effects, standard deviations and variance-covariance matrices. These posterior estimates were used for general linear hypothesis testing of two types of contrasts: deviation contrasts of one population compared to the mean of the other populations, and pairwise contrasts between all pairs of populations.

#### Pi1

To assess replication of eQTLs and sQTLs across populations, we calculated pi1^34^ by the method of Storey, using the π_0_ statistic from the qvalue package (π_1_ = 1 - π_0_) to compare eQTLs discovered in both populations. For eQTLs, we used the top 3000 QTLs (best SNP-eGene pair) in any one population across the other populations. For this analysis, we excluded YRI due to the smaller sample size and subsequently reduced power to detect and replicate QTLs. To assess replication of the AFGR population eQTLs in another continental group we also tested replication of eQTLs in GEUVADIS European population bulk-called eQTLs. We tested replication only when the best SNP for a SNP-eGene pair was present in both populations (**Supplementary Figure 3**).

For the set of significant SV-gene pairs (10% FDR) in one population, we calculated π_1_ on their nominal pass p-values in the other population. We determined 95% confidence intervals for these π_1_ estimates with bootstrapping. To stratify π_1_ on the frequency differences of SVs between the African and Geuvadis dataset, we calculated Wright’s *F_ST_* between the African and European Geuvadis samples with plink^93^ v1.90b6.4.

#### Allelic differentiation

For each top eQTL variant identified by meta-analysis, allele frequency was calculated from the African (ESN, GWD, LWK, MSL, YRI) populations in the 1K Genomes Project and MKK, and European (CEU, FIN, GBR, TSI) populations in 1K Genomes Project. When the meta-analysis variant was specific to MKK (ie, not present in the 1K Genomes VCF), these variants were omitted from further analysis (266 (3%) eQTL variants and 327 (4.7%) sQTL variants). When the meta-analysis variant was not genotyped in MKK, allele frequency in the African populations was calculated using the 1K Genomes populations only. The absolute value of difference in allelic frequency in African and European populations was used to identify variants with high differential allele frequencies and their associated genes for enrichment analysis.

#### Calculation of Fst and iHS

We calculated Weir and Cockerham’s Fst^37^ using vcftools^94^ v0.1.16 for each population pair, on a per-site basis. At each site, for each focal population, an average Fst value was calculated across the four population comparisons. These average values were sorted for each focal population, and we subsequently determined the SNPs that comprised the top 1% of average Fst values.

iHS was calculated using selscan^95^ v1.2.0. We first used vcftools to generate files in IMPUTE format (using --IMPUTE), then transposed rows and columns to produce the haplotype input for selscan. We generated map files for input to selscan assuming 1 cM/Mb. We ran selscan with default settings. In order to avoid issues concerning accuracy of SNP polarization, and to allow for detection of possible selection on ancestral alleles, sites were coded as 0 and 1 according to reference/alternative rather than ancestral/derived, and iHS values that were extreme in either the negative or positive direction were considered of interest. We standardized iHS scores using selscan, in 100 allele frequency bins. We focused on sites with extreme iHS scores, defined as an absolute value greater than 4.

To identify eQTL and sQTL signals with evidence of selection, we intersected the list of variants with high iHS (absolute value > 2) and high Fst (top 1% per population) with lead QTL variants and variants in high LD with lead QTL variants (r2 > 0.8).

### Allele Specific Expression (ASE) from RNA-seq

To assess allele specific expression, we identified heterozygous variants in each individual, and confirmed these sites using the high-coverage (30x) genomes from the New York Genome Center (http://ftp.1000genomes.ebi.ac.uk/vol1/ftp/data_collections/1000G_2504_high_coverage/working/20190425_NYGC_GATK/, accessed 11 May 2019). On average, 93% of the heterozygous sites in the hg38-aligned VCFs from 1K Genomes were also heterozygous in the high-coverage genomes and we retained these sites for ASE analysis.

FASTQs were recreated from aligned-read BAM files using the samToFastq tool in Picard, with default settings and excluding non-primary alignments. Allele specific expression was measured using WASP^96^ to account for reference mapping bias using the workflow at (https://github.com/durrantmm/ase_read_counting_workflow), with the addition of a mapping quality filter > 10 and base quality filter > 2 to ASEReadCounter in GATK^97^. Heterozygous sites passing these quality filters and with at least 10 reads in an individual were tested for ASE using a beta binomial distribution.

Sites with allelic ratios between 0.05-0.35 or 0.65-0.95, and passing false discovery error rate of < 0.05 in any individual were considered to exhibit ASE. As a control, we examined the number of sites and genes tested for ASE and the number of sites and genes displaying significant ASE across individuals (aseGenes; **Supplementary Figure 5**).

#### Protein truncating variant annotation

We used Variant Effect Predictor^98^ (Ensembl 90/GENCODE v27) with the LOFTEE^99^ plugin (v1.0.4) to identify protein truncating variants (PTVs, a.k.a. nonsense variants) expected to result in loss of function in all variants in our study. PTVs were defined as any variant predicted by VEP to gain a premature stop codon. We restricted our analysis set to high confidence PTVs based on the “LoF” annotation from LOFTEE that also contained no warnings in the “flag” column. This yielded 8,360 total high confidence PTVs across all AFGR samples, 342 of which had sufficient RNA-seq coverage (at least 10 total reads) for testing allelic expression.

Because most protein-truncated variants are expected to undergo nonsense-mediated decay and thus may have a biologically-founded prior for an extremely high or low reference ratio, we extended this analysis to include variants with an ASE reference ratio >0.95 or < 0.05. No predicted PTVs showed evidence of extreme alternate-allele-biased expression (reference ratio < 0.05), but 59 presented with extreme reference-allele-biased expression (reference ratio > 0.95), consistent with expectations of ASE driven by nonsense-mediated decay.

To determine if any PTVs without allele-specific expression (ie, escaping nonsense-mediated decay) fell within genes with predicted PTV-induced gain-of-function effects, we took the union of genes reported by Coban-Akdemir *et al* (2013)^42^ in table S1 (OMIM genes with reported NMD escape intolerance), table S3 (genes depleted for truncating variants in both the ARIC and ExAC databases), and table S4 (genes depleted for truncating variants in either the ARIC or ExAC databases). We further annotated genes with disease associations from the Online Mendelian Inheritence in Man (OMIM) database (morbidmap.txt, accessed 22 Apr 2022)^100^. The OMIM gene to phenotype map was filtered to gene-phenotype associations with a minimum phenotype mapping key value of 3 (indicating a known molecular basis for the disorder implicating the gene), and reformatted to produce a file with a single entry summarizing all associated diseases per gene, which is available on our github (see **Code**). Gene names from all annotation sources were mapped to the current hgnc gene symbol then converted to ensembl IDs using the ensembl BioMaRt^101^ before merging to our ASE results based on ensembl ID match. A gene was considered a gain-of-function gene if it appeared in the union set of PTV-depleted genes, and an OMIM disease gene if it had at least one confidently associated disorder.

#### Identification of low-frequency allelic effects

We defined low-frequency ASE variants as those with a minor allele frequency in AFGR < 0.05, our minimum allele frequency threshold for QTL calling. Variants with significant evidence of ASE in at least one sample were additionally annotated with the corresponding allele frequency calculated from the 30x coverage VCFs of the 1000 Genomes European populations: Utah residents (CEPH) with Northern and Western European ancestry (CEU), British From England and Scotland (GBR), Finnish in Finland (FIN), Toscani in Italia (TSI). There were no flipped alleles between the VCFs. Variants with a MAF of 0.0 in high-coverage European population genotypes are described as monoallelic and variants with a missing allele frequency are described as not annotated. The majority of the non-annotated variants 10245/10246 were MKK-specific variants and are likely due to the MKK samples undergoing sequencing the variant calling separately from the 1000 Genomes samples. The remaining missing variant, 3_104345691_T_C, demonstrated ASE in a single YRI sample; there is a short indel just upstream (chr3_104345688_TTA_T) that seems to be homozygous variant in EUR populations that may be affecting ability to determine genotype at 3_104345691_T_C.

#### Identification of eQTL-independent allelic expression events

To identify rare allelic effects that are not tagging a more common eQTL-associated variant, we identified ASE events that are observed without a heterozygous eSNP for the given ase/eGene in the ASE-presenting samples. Nominally significant individual-level ASE and meta-analyzed eQTL results were joined on the reported ase/eGene using bash join. For a sample demonstrating ASE for a given gene, the corresponding genotype of each eSNP associated with that gene was extracted from the merged AFGR VCF. Individual-level ASE variants were subsequently filtered out if the encompassing aseGene had one or more associated eSNPs for which the individual was also heterozygous. Remaining variants that were significant for ASE in at least one individual who did not harbor a heterozygous eQTL for the same gene were deemed “eQTL-independent ASE events.”

As a side note, MKK sample IDs have a different encoding in the merged VCF than in other AFGR datasets, so an additional sample ID map was used to link MKK sample IDs with the MKK VCF IDs (available on the github).

### Allele-Specific Chromatin Accessibility (ASC) from ATAC-Seq

Similar to ASE from RNA-seq, we confirmed the presence of heterozygous sites in high coverage (30x) genomes prior to testing allele-specific chromatin accessibility (ASC). Allele-specific chromatin accessibility was measured using WASP^96^ to account for reference mapping bias using the workflow at (https://github.com/durrantmm/ase_read_counting_workflow), with the addition of a mapping quality filter > 10 and base quality filter > 2 to ASEReadCounter in GATK. Heterozygous sites passing these quality filters and with at least 10 reads in an individual were tested for ASC using a beta binomial distribution. Sites tested in at least 5 individuals, with allelic ratios between 0.05-0.35 or 0.65-0.95, and passing FDR < 0.05 in at least 50% of individuals tested were considered to be sites exhibiting ASC across all populations. We investigated the distribution of the number of sites and genes tested across individuals, as well as the number of sites and genes displaying significant ASC across individuals (**Supplementary Figure 5**).

#### ASC sites in Activity by Contact (ABC) enhancers

Activity by Contact (ABC) enhancer predictions connect enhancers to genes in a cell-type-specific context by integrating an enhancer’s estimated activity strength, the frequency of 3D contact between the enhancer and the promoter of the target gene, and the estimated impact of the the enhancer’s inhibition on the gene expression based on CRISPR-inhibition assays^102^. We accessed information on enhancer-gene relationships for B-cell biosamples or GM12878, an LCL biosample from https://www.engreitzlab.org/resources (accessed May 16 2023)^47^. Significant allele-specific chromatin accessibility (ASC) sites identified in at least one individual were intersected with these ABC predictions using `bedtools intersect -wao`, which required that the ASC variant rest within the enhancer element.

#### High-confidence ASC sites

To identify ASC sites robust to experimental noise and cell state, we compared sites across the whole group to identify a set of 7,559 high-confidence ASC sites observed consistently (defined here as tested in at least 5 individuals and significant in more than 50% of those tests with the same direction of effect in all significant samples) across AFGR.

### Machine Learning Models Predicting Chromatin Accessibility

#### Data preparation

ATAC-seq reads for each individual were mapped to GRCh38/hg38 reference genomes using Bowtie2^103^ aligner. Pair-end reads that were mapped to the same genomic positions were deduplicated. We selected six samples with the highest coverage for each ancestry and merged the filtered BAM files to generate ancestry-specific BAMs. We performed a +4/-4 shift on the ancestry-specific BAMs and then generated BigWig tracks for each ancestry using bedtools genomecov -5 -bg followed by bedGraph to BigWig conversion using bedGraphToBigWig^104^. Ancestry-specific peaks were called using ENCODE ATAC-seq pipeline (http://doi.org/10.5281/zenodo.3662027) for all six ancetries.

#### Model training

ChromBPNet is a base-resolution convolutional neural network model of chromatin accessibility and performs automatic Tn5 bias correction by taking a pre-trained bias model to regress out the enzyme biases. We used the bias model from the ChromBPNet repository (https://zenodo.org/record/8011885) and trained six ChromBPNet models, one for each of the six ancestral populations, using the 5 samples with the deepest ATAC-seq sequencing per group. Each ChromBPNet model takes a 2114-bp sequence around the ATAC-seq peak summit and outputs the base pair resolution profile and the total number of Tn5 insertions in the center 1000 base pairs for its ancestry. We augmented the datasets by incorporating the reverse complements of the sequences and added random shifts (up to 500 bp) to the input sequences. Each model was trained using five-fold cross-validation, where different chromosomes were held out for testing and validation during each fold. Our chromBPnet model provides predicted ATAC-Seq counts at each allele of a SNP, which allows us to calculate a log fold-change (logFC) prediction for chromatin accessibility at any position of the genome, even in the absence of the heterozygosity required for the empirical determination of ASC. The logFC prediction values were well-correlated between populations (**Supplementary Figure S19**), and were subsequently averaged to combine the models into a single annotation per variant.

#### Calculating chromBPnet prediction significance

For each chromosome, we sampled 1 million variants from the list falling in that chromosome. We shuffled the 2114 bp sequence around each SNP, centered at the SNP, while preserving dinucleotide frequency, made 2 copies of each shuffled sequence, then inserted each allele of the SNP at the center of the shuffled sequence to create a set of null SNPs. We scored each of these null SNPs with the model in the same way that we scored each observed SNP. Then, for each observed SNP, we calculated what proportion of the null SNPs have an equally high or higher (more extreme) score, to generate an empirical p-value. When averaging scores across folds and ancestries, we also calculated the average for the null scores at each SNP across folds and ancestries and calculated the empirical p-value for the mean of the observed scores with respect to the distribution of the mean null scores across folds and ancestries for all 22 chromosomes (22 million null variants).

#### Motif Identification and deepSHAP Model Interpretation

We utilized TF-MoDISco^49^ on DeepShap^61^ interpretations derived from 30,000 randomly selected ATAC-seq peak regions to identify motifs learned by each of the six ancestry-specific models. As expected, TF-MoDISco generated almost identical sets of motifs for all six models. Subsequently, we annotated 77,663 variants that are significant (LFC p-value < 0.001) for the motif sites surrounding the variants. We computed the DeepLift^50^ interpretation for both the reference and alternative alleles using the 2114-bp sequence around the variants and averaged across the six ancestry-specific models. We then utilized TomTom^51^ to search for matches to the TF-MoDISco seqlets within the variant-centered 30bp windows. We subsequently filtered to human transcription factor motifs and applied a TF-MoDISco q-value threshold of 0.05 to identify variants with significant impacts on transcription factor binding motifs. A motif was defined as “destroyed” if the reference allele produced a significant motif while the alternate allele did not, “created” if the alternate allele produced a significant motif while the reference allele did not, and “swapped” if both the reference and alternate alleles contributed significantly to binding motifs for different transcription factors.

### Colocalization of GWAS and QTL signals

#### GWAS Selection

We integrated the AFGR caQTLs, meta-analyzed AFGR eQTLs, and meta-analyzed European GEUVADIS eQTLs with large, publicly available GWAS across a range of traits and diseases. Subsequent LD-score regression, colocalization, and fine-mapping analyses were performed on a curated list of GWAS summary statistics, including blood biomarkers (e.g. lymphocyte count), common complex diseases (e.g. Type 2 diabetes), and conditions with etiology related to immune function (e.g. Crohn’s disease). A GWAS region was defined as a significant GWAS variant (p-value <= 5e-8) at least 1Mb away from another GWAS signal. A full list of GWAS, corresponding publications, and analysis results can be found in **Supplementary Table 9**.

#### LD Score Regression

Heritability estimation and partitioning were performed by LDSC (or S-LDSC) [https://github.com/bulik/ldsc]^105,106^. For this analysis, we computed LD scores (--l2) in a set of 358 individuals of European ancestry from GEUVADIS as a proxy LD panel for the predominantly European GWAS. We then partitioned heritability (--h2 --annot) using a set of 98 variant annotations: the 97 “baseline” annotations from the S-LDSC publication^105^, plus an annotation for whether variants lie in an ATAC seq peak (from either GM12878 [GEO accession ID: GSE47743], or a unified set of narrow peaks derived from our data; see “ATAC-Seq Library Preparation and Sequencing”).

#### GWAS-QTL Colocalization

GWAS associations were subjected to a two step process to first identify regions overlapping QTL variants, then to test for evidence of shared genetic architecture between the QTL and GWAS signals using coloc^31^. GWAS associations that met a genome-wide significance threshold of 5×10-8 were selected as candidate regions for downstream analysis. Lead SNPs were ranked by significance, and the most significant SNPs that were more than 1Mb apart were then assessed for nearby QTL variants. The 1Mb buffer was selected to ensure that no variants were double-tested for colocalization downstream. Subsequently, a GWAS region was prioritized for colocalization if there was at least one nominally significant QTL variant (p <= 1e-5) within 10kb of the sentinel GWAS SNP. Linkage disequilibrium (LD) blocks range from <1KB to >100kb, with averages around 10kb^107^. Thus, GWAS regions prioritized for colocalization analysis were putatively independent signals, containing a genome-wide significant variant that was within 10kb of a significant AFGR eQTL, AFGR sQTL, AFGR caQTL, or European GEUVADIS eQTL variant.

To identify which of these regions likely shared underlying genetic architecture with any of the nearby QTL signals, all variants within 500kb upstream and downstream of the sentinel GWAS variant and present in both the GWAS and the QTL dataset were tested for colocalization with *coloc* v5.1.0.1^31^. A GWAS region was tested for colocalization with all significant QTL signals within 10kb of the sentinel variant, so a given GWAS region could have been tested for shared genetic architecture with multiple eGenes, splice junctions, and open chromatin regions. Moderate colocalization was deemed as a posterior probability of shared genetic architecture (COLOC_h4) >= 0.5, while strong colocalization was defined as a posterior probability >= 0.8.

It is worth noting that coloc, as implemented here, assumes datasets are derived from populations with identical patterns of LD, an assumption that may not hold in the context of European GWAS and African QTLs. Additionally, this implementation of coloc assumes there is at most one causal variant at the locus.

#### Fine-mapping of GWAS with QTL signals

Colocalized regions were fine-mapped with the corresponding QTL and GWAS summary statistics independently using *SusieR*^108^, and assuming one causal signal. SusieR is a Bayesian statistical method that leverages summary statistics and linkage disequilibrium information to construct credible sets and prioritize likely causal variants. The GWAS variants were filtered to those present in the European GEUVADIS VCF, which was used to calculate a proxy LD panel for each colocalized GWAS region. For each region, QTL summary statistics were subset to variants present in the filtered GWAS dataset, and an in-sample LD panel was constructed from the VCF corresponding to the QTL set selected. For the GWAS and QTLs, variant-level Z-scores were calculated from the beta and standard errors according to equation (6) from Zou *et al* (2022)^26^. Other than the maximum number of independent signals, `susie_rss` was run with default parameters.

## Data Availability

The raw sequencing data generated for this project are available at ENCODE: https://www.encodeproject.org/search/?searchTerm=AFGR&type=Experiment

The QTL summary statistics and meta-analyses are available at: https://github.com/smontgomlab/AFGR

Previously published databases or datasets used in this work

- 1000 Genomes genotyping aligned to GRCh38 (http://ftp.1000genomes.ebi.ac.uk/vol1/ftp/data_collections/1000_genomes_project/release/20190312_biallelic_SNV_and_INDEL/)
- 1000 Genomes genotyping 30x coverage (http://ftp.1000genomes.ebi.ac.uk/vol1/ftp/data_collections/1000G_2504_high_coverage/working/20190425_NYGC_GATK/)
- GEUVADIS eQTLs (ftp://ftp.ebi.ac.uk/pub/databases/microarray/data/experiment/GEUV/E-GEUV-1/analysis_results/)
- Genome Tissue Expression database (GTEx) (https://gtexportal.org/home/)
- gnomAD (https://gnomad.broadinstitute.org/downloads)
- African Pangenomic Contigs (contigs: https://www.ncbi.nlm.nih.gov/Traces/wgs/PDBU01?display=contigs; contig annotations: https://www.nature.com/articles/s41588-018-0273-y#Sec18 Tables S1-4)
- Itan Lab’s GOF/LOF database (https://itanlab.shinyapps.io/goflof/)
- ClinGen Dosage Sensitivity database (https://search.clinicalgenome.org/kb/gene-dosage)
- Collins et al, dosage sensitivity resource (https://www.sciencedirect.com/science/article/pii/S0092867422007887?via%3Dihub#app2 Table S3)
- OMIM (https://www.omim.org/)
- Open Targets (https://www.opentargets.org/)

## Code Availability

The code for generating and analyzing the AFGR data will be available at: https://github.com/smontgomlab/AFGR

## Acknowledgements

This work was supported in part by the National Institutes of Health through ENCODE4 U01HG009431 (MKD, PCG, AK and SBM), R35GM133747 (RCM) and 1F31HG012495 (SMY). MA was supported by the National Library of Medicine under training grant T15LM007033. NH was supported by an NSF GRFP.

## Author Contributions

MKD, PCG, JF, PJM, BH, RCM, AK, MSS, DG and SBM contributed to the conception of this work. MKD, PCG, EK, JF, PJL, RCM, MSS, DG and SBM contributed to the study design. MKD, CP, KSS contributed to raw data acquisition. MKD, PCG, EK, SK, SMY, DN, MA, TC, ZC, VRD, MPSM, CP, CJS, KSS, LAS, DG generated resource data. MKD, PCG, SK, SMY, DN, NA, MA, TC, NH, KBR, CJS, RAU, BB, LAS, DG contributed to data analysis. MKD, PCG, SK, SMY, DN, NA, MA, MJG, KBR, RAU, BB, JF, PF, LAS, MSS, DG and SBM contributed to data interpretation. MGD, MJG contributed to software development. MKD, PCG and SBM wrote the manuscript.

---

**Extended Data Figure 1.**
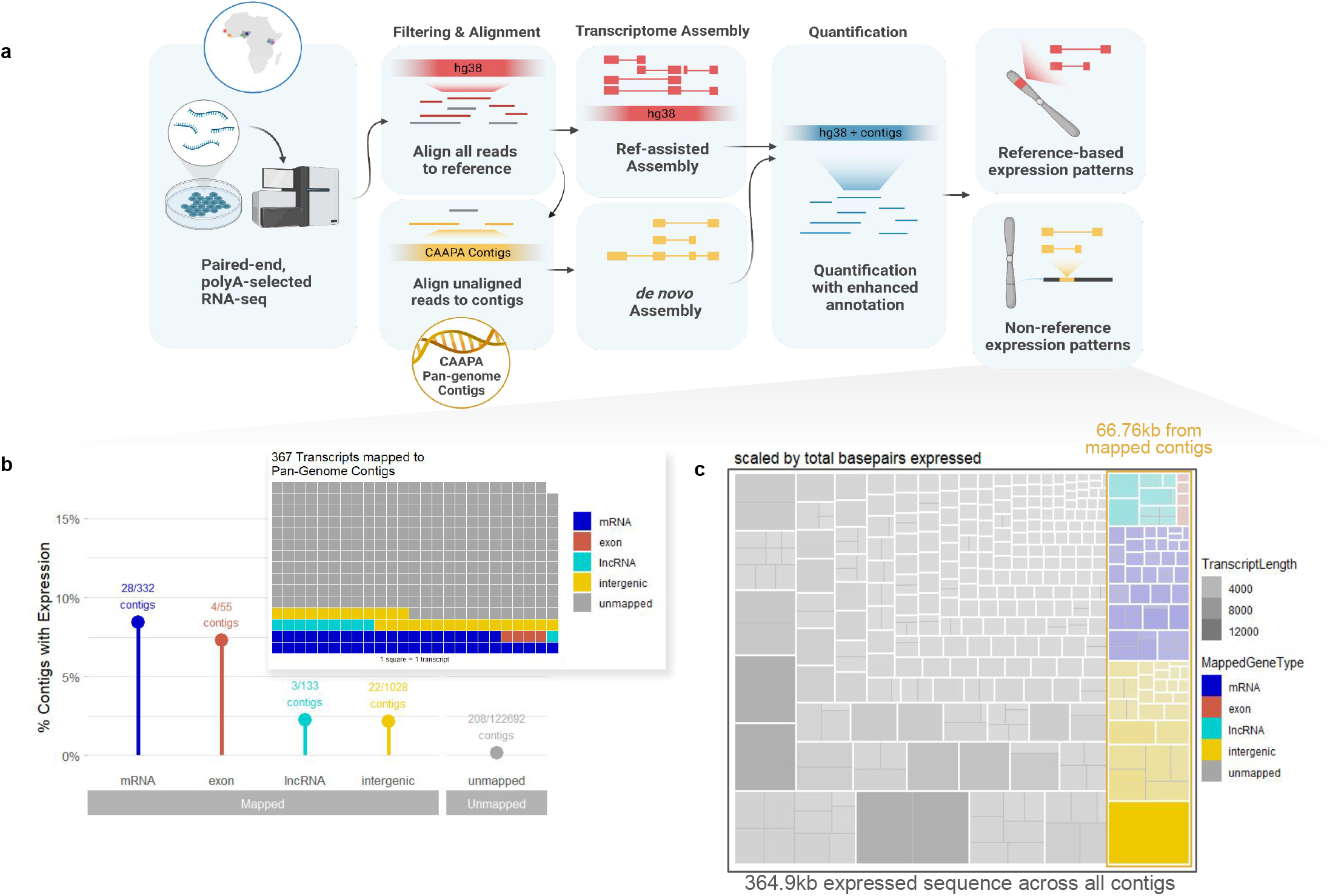
Expression of novel sequences transcribed from genome contigs missing from GRCh38. a. Pipeline for alignment and identifying novel reference genome and non-reference genome transcripts. b. The proportion of contigs that are anchored in each genomic region type, or remain unmapped, that harbor at least one expressed transcript over the total number of contigs anchored to that region type. Inset summarizes the mapping status of the parent contigs for the 367 transcripts originating from the non-reference genetic sequences; each square is a transcript. c. Treeplot of transcripts that mapped preferentially to the Sherman at al^2^ CAAPA contigs. Each box outlined in white represents a separate contig with at least one transcript; colored boxes indicate what type of region Sherman et al mapped the contig within and grey boxes indicate the contig was not anchored in the reference genome. The size represents the total number of expressed bases originating from that contig. Within each contig, boxes delineated with grey lines represent individual transcripts mapping to the contig, where the relative size and opacity represent the transcript length (in bp).

**Extended Data Figure 2:**
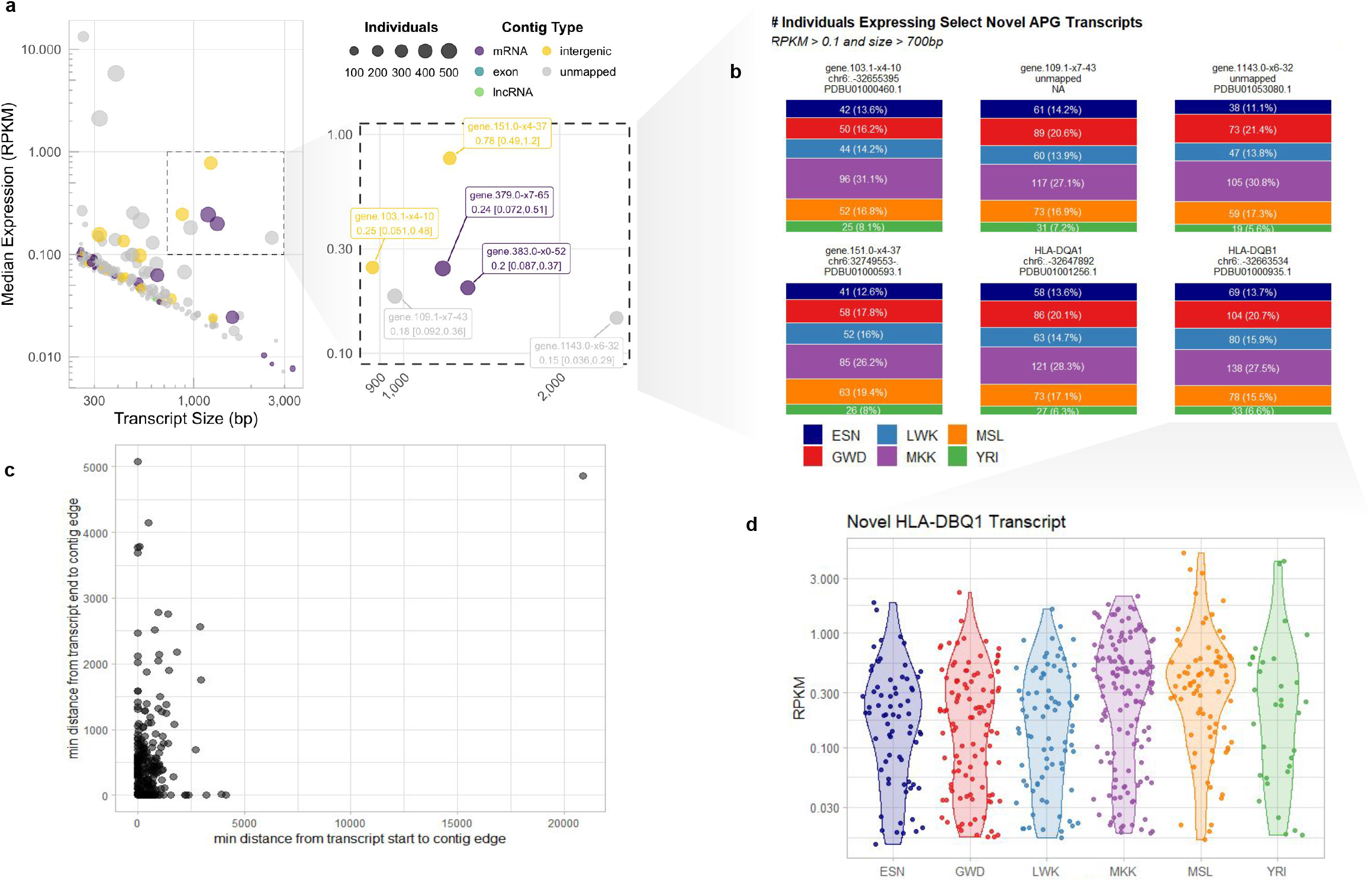
Characterization of expressed sequences transcribed from genome contigs missing from GRCh38. a. Scatter plot showing transcript size vs expression of transcripts mapped to the CAAPA contigs. Each point is a transcript, scaled by the number of individuals it was detected in, and colored by the context of the contig it mapped to (mRNA, exon, lncRNA, intergenic, or unmapped in Sherman et al.). Inset highlights six genes with median RPKM > 0.1 and transcript length > 700 bp. b. Stacked bar plots depicting the number of individuals with expression of the six transcripts highlighted in (E) across all AFGR populations. c. Scatter plot showing the minimum distance from the start of the transcript (Y-axis) and the end of the transcript (X-axis) to the edge of the contig. d. Gene expression distributions for a novel transcript of the HL-DQB1-mapping contig are similar across populations.

**Extended Data Figure 3:**
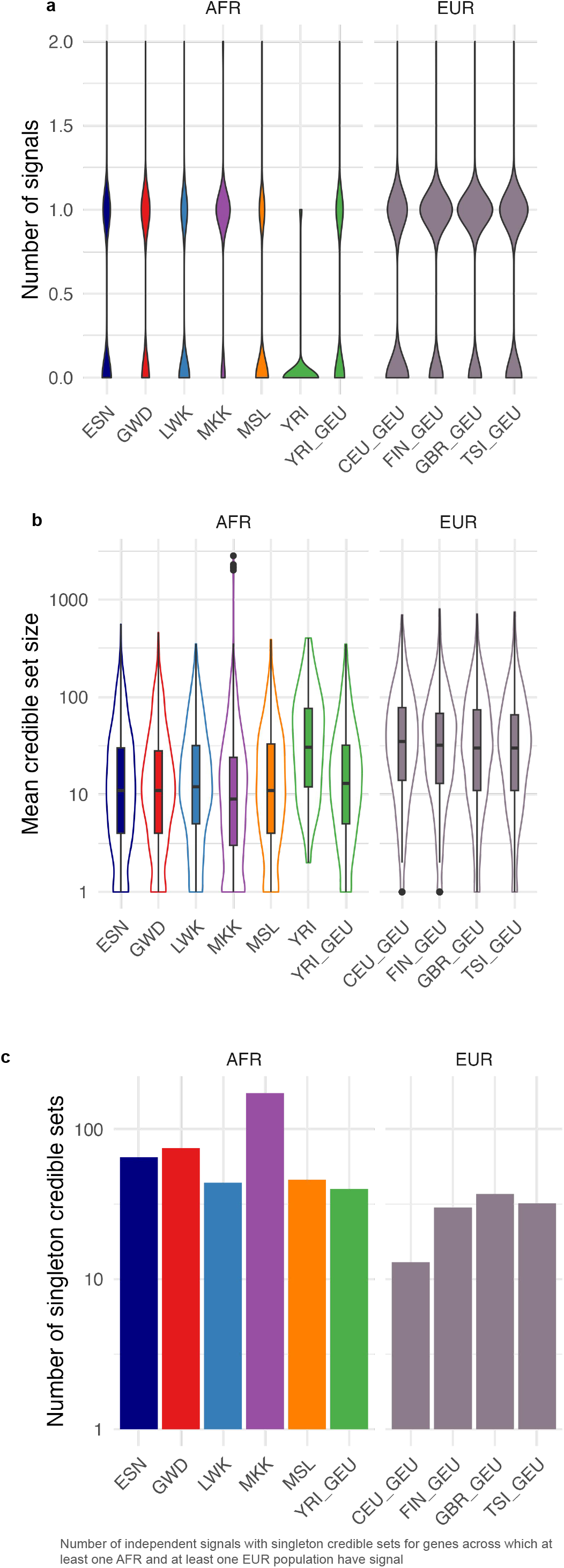
QTL-fine-mapping and credible set sizes. a. Number of independent signals detected by SusieR, per eGene, for all genes with a credible set in at least one African and at least one European population. b. Average credible set size per fine-mapped eGene across populations, for all genes with a credible set in at least one African and at least one European population. c. Number of credible sets with a single variant in the given population across all genes with a credible set in at least one African and at least one European population.

**Extended Data Figure 4:**
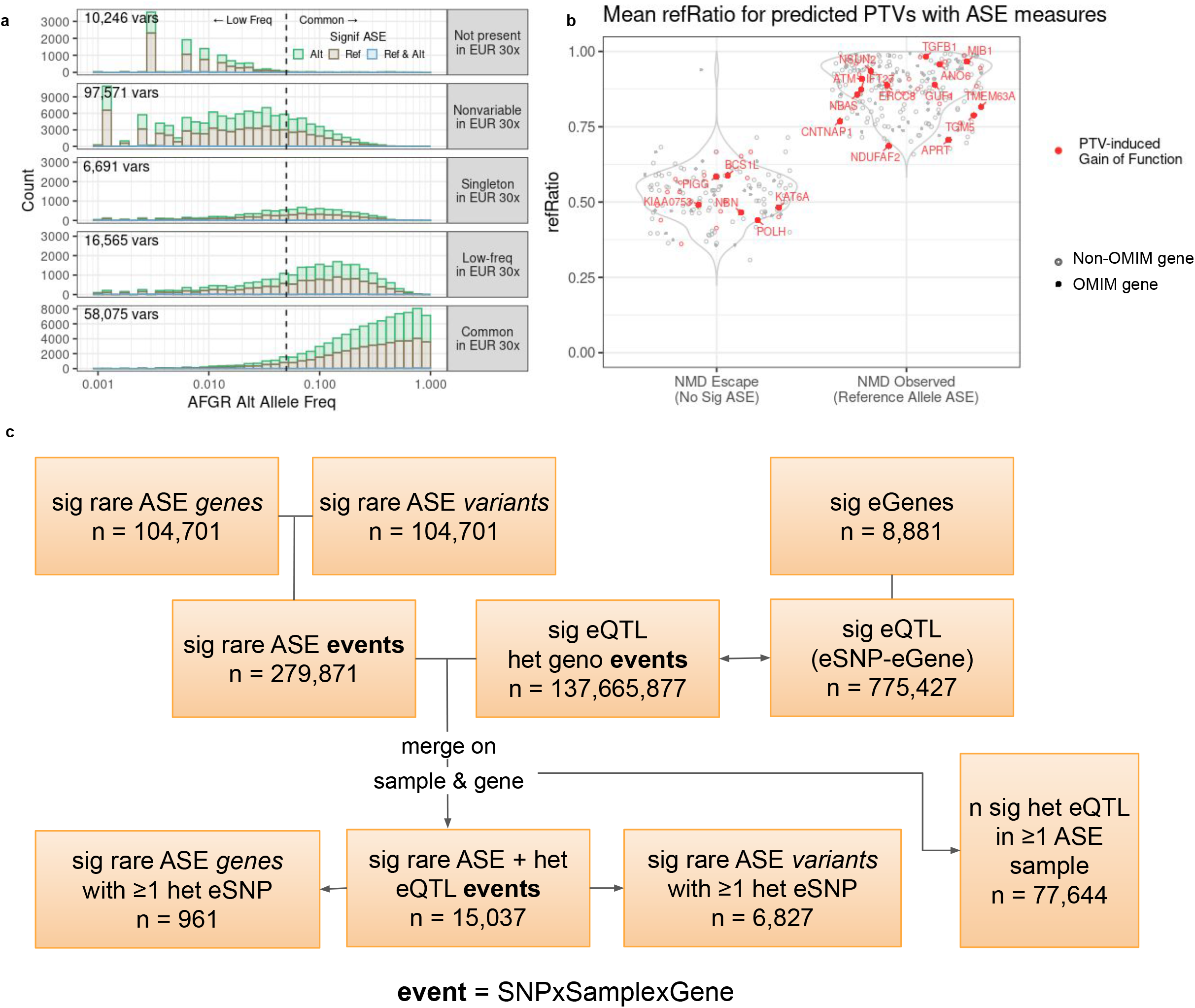
Trends in variants exhibiting allele-specific expression. a. Allele frequency spectra in African populations for variants displaying significant allele-specific expression (ASE), grouped by allele frequency status in the 1000 Genomes European populations. b. Average reference ratio measures for rare protein-truncating variants tested for ASE in AFGR. Variants in red are within genes with predicted PTV-induced gain-of-function effects. Solid points are known OMIM genes; variants in known OMIM genes with predicted PTV-GoF effects are labeled with the encompassing gene name. c. Flowchart illustrating the intersection of rare (MAF < 0.05) ASE variants with heterozygous eQTL variants.

**Extended Data Figure 5:**
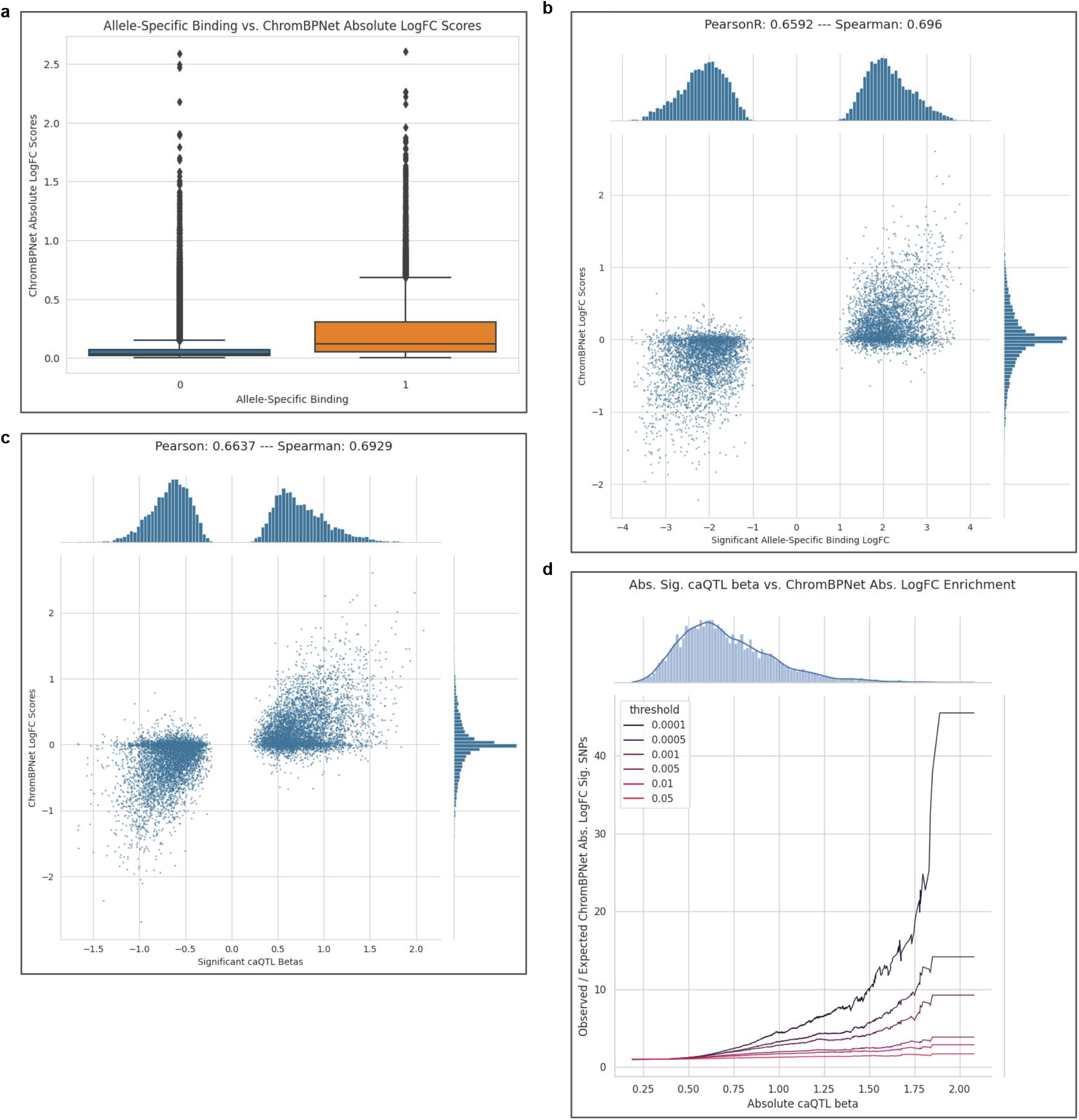
ChromBPnet quality control. a. ChromBPnet logFC predictions for chromatin accessibility at sites tested for ASC, stratified by whether the site was determined to exhibit significant ASC in at least one sample (Allele-specific chromatin accessibility = 1) or not (Allele-specific chromatin accessibility = 0). b. Correlation of chromBPnet-predicted chromatin accessibility logFC scores and the average observed logFC in chromatin accessibility across samples exhibiting significant ASC (n=7,559). c. Correlation of chromBPnet-predicted logFC scores and caQTL effect sizes (betas) at significant caQTL variants (n=11,098). d. Enrichment of absolute caQTL effect size with absolute predicted allelic logFC at increasingly stringent chromBPnet permutation p-value thresholds.

**Extended Data Figure 6:**
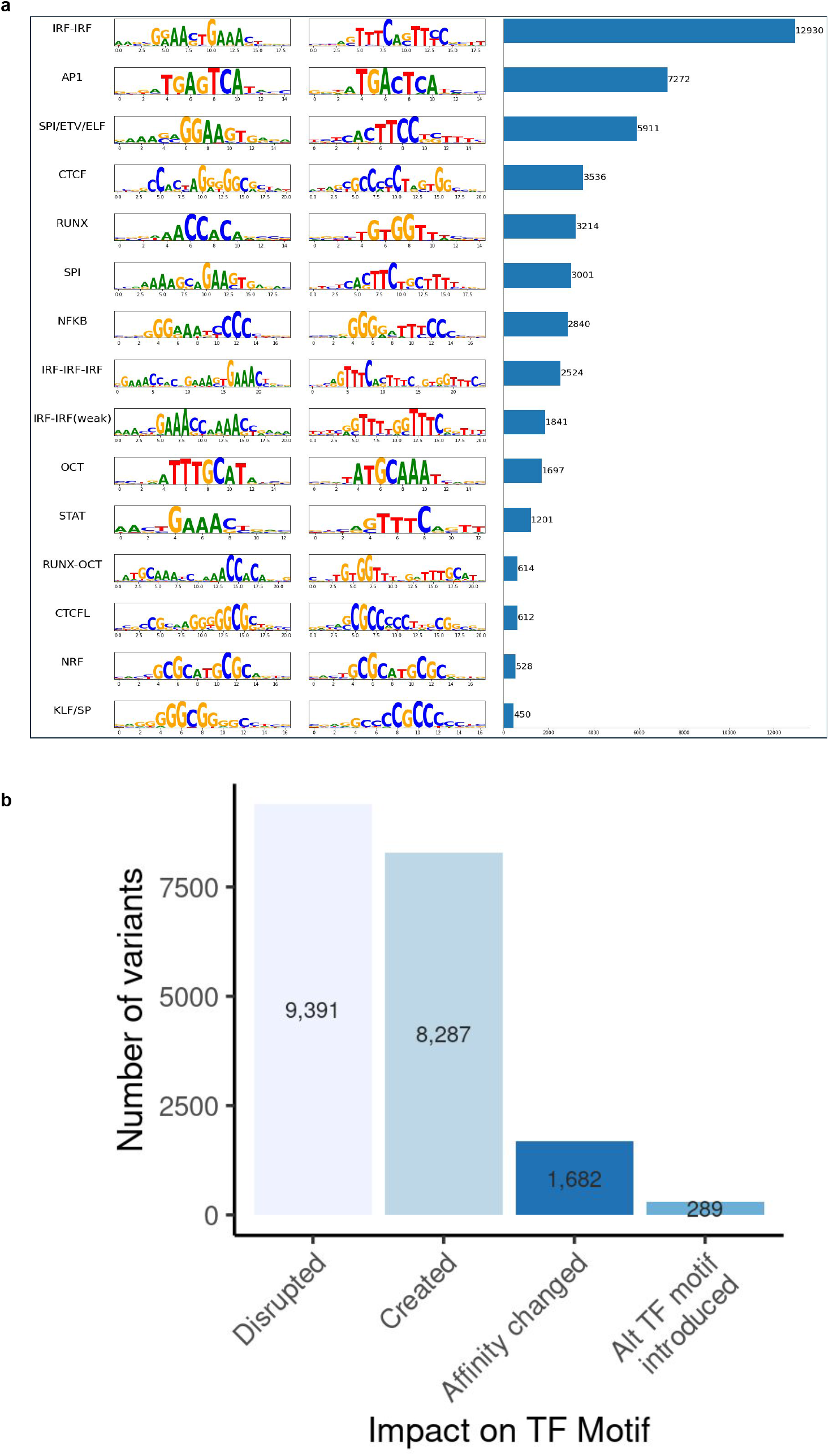
Predicted impact on transcription factor binding sites. a. Top 15 most frequent transcription factor binding sites motifs predicted to be impacted by variants using chromBPNet score (logFC p-values < 0.001) and DeepShAP. b. Frequency of predicted impacts on human transcription factor binding motifs.

**Extended Data Figure 7:**
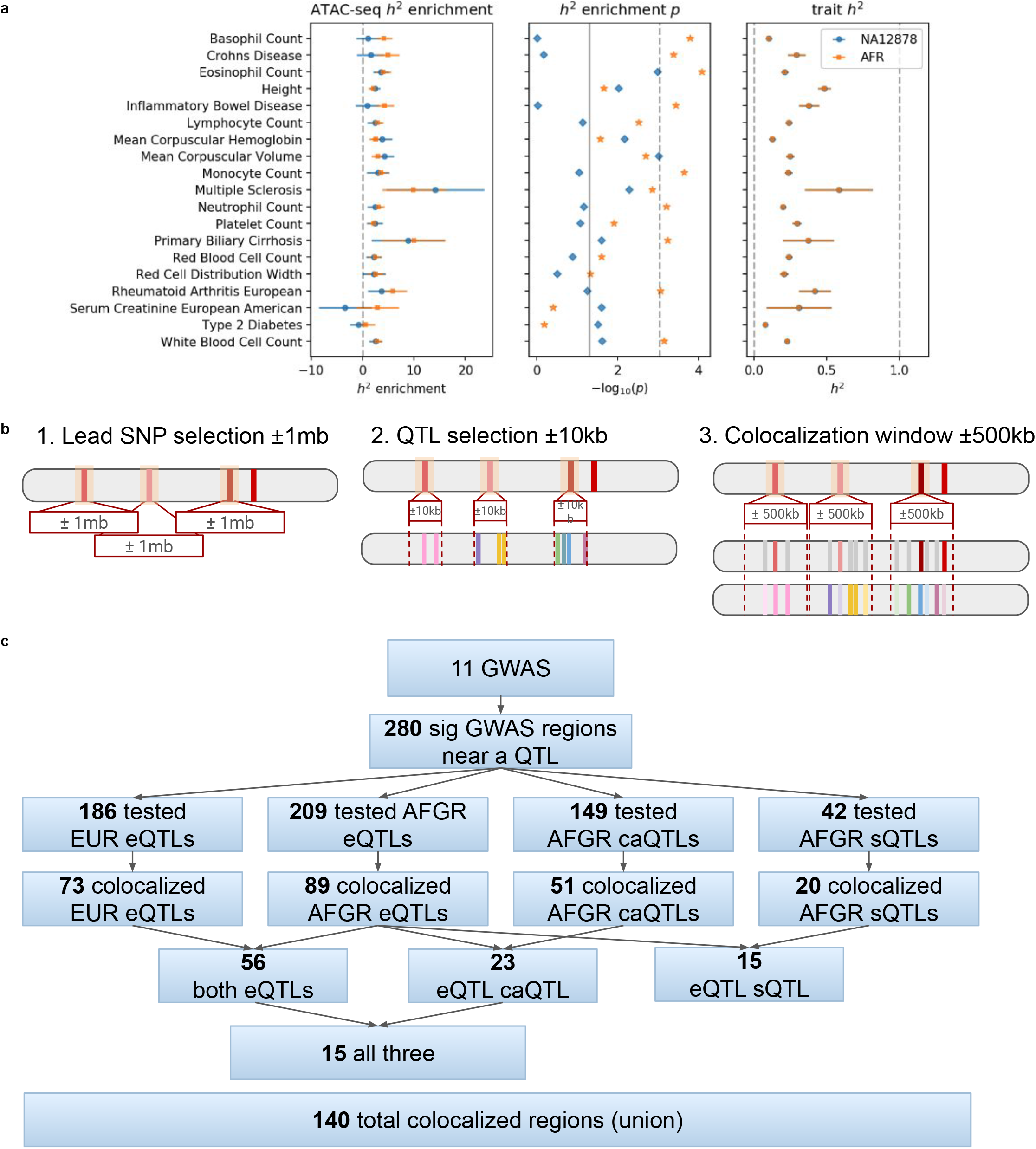
LD Score Regression & colocalization region selection. a. LD-Score enrichment values for selected GWAS. b. Schematic depicting prioritization of regions and variants used in colocalization testing. c. Flow chart summarizing number of significant GWAS regions identified, prioritized for colocalization with each QTL type, and colocalized with posterior probability of a shared causal signal >= 0.5

**Extended Data Figure 8:**
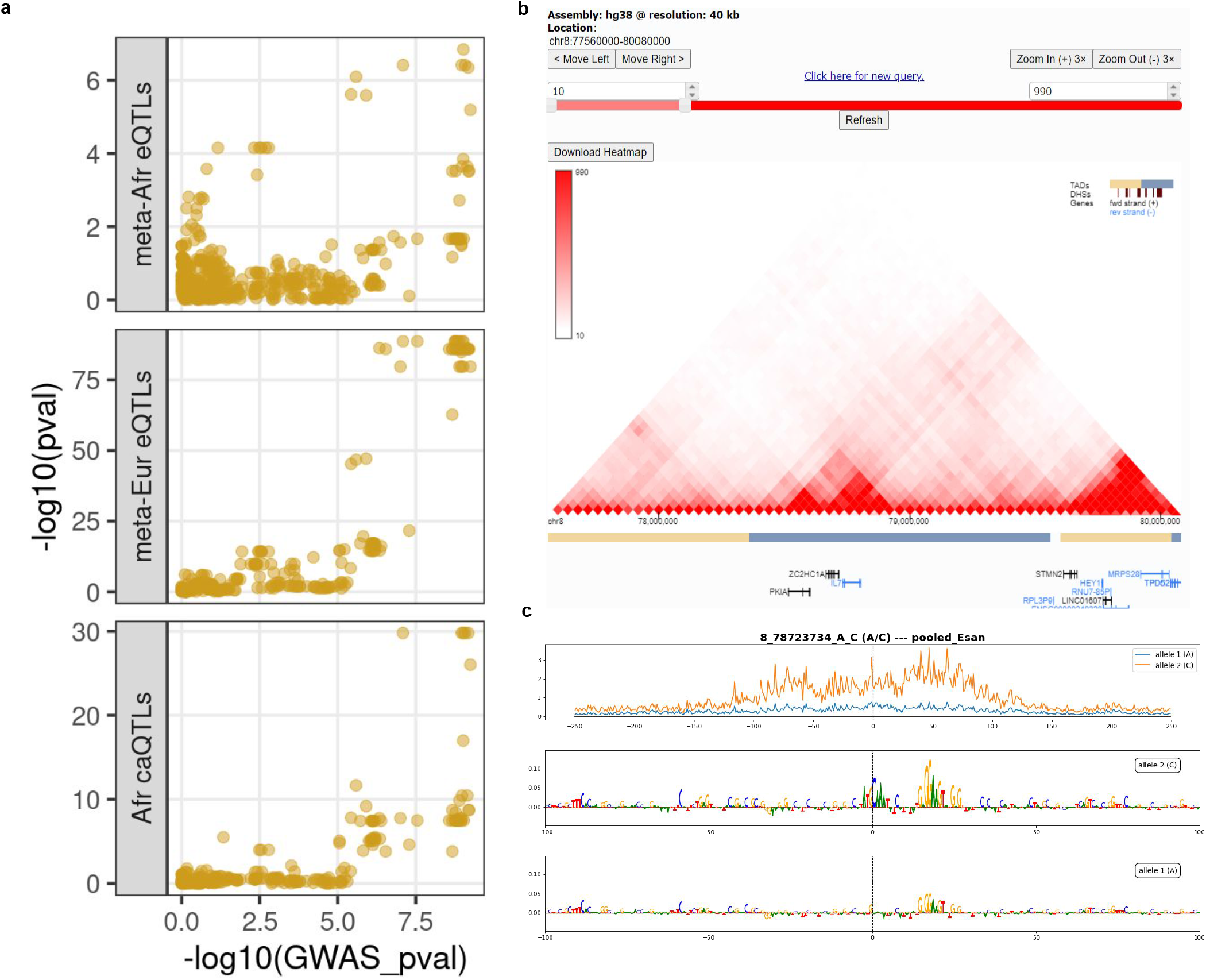
IL7 locus additional figures. a. Locuscompare plots at the IL7 locus showing the correlations for the meta-African IL7 eQTL p-values vs multiple sclerosis GWAS p-values (top); meta-European IL7 eQTL p-values vs multiple sclerosis GWAS p-values (middle); African caQTL p-values for one of three open chromatin regions at the TES of IL7 vs multiple sclerosis GWAS p-values (bottom) b. Hi-C data from 3DGenome showing interaction between IL7 TES and TSS c. [top] Predicted chromatin accessibility of the C allele (orange) and A allele (blue). [bottom] deepSHAP scores across IL7 locus in the presence of the C and A alleles, respectively.

**Extended Data Table 1:** QTL statistics.

| Population | QTL Type | Number of significant QTL features (genes, splice clusters or open chromatin regions) |
| --- | --- | --- |
| ESN | eQTL | 2,205 |
| GWD | eQTL | 2,601 |
| LWK | eQTL | 1,532 |
| MKK | eQTL | 3,923 |
| MSL | eQTL | 1,291 |
| YRI | eQTL | 150 |
| AFGR | SV-eQTL | 2,611 |
| AFGR | meta eQTL | 8,881 |
| GEU | meta eQTL | 6,767 |
| ESN | sQTL | 574 |
| GWD | sQTL | 660 |
| LWK | sQTL | 505 |
| MKK | sQTL | 842 |
| MSL | sQTL | 473 |
| YRI | sQTL | 142 |
| AFGR | meta sQTL | 6,949 |
| AFGR | caQTL | 10,569 |

