## Supplementary Figures for "Transcriptomics and chromatin accessibility in multiple African population samples"

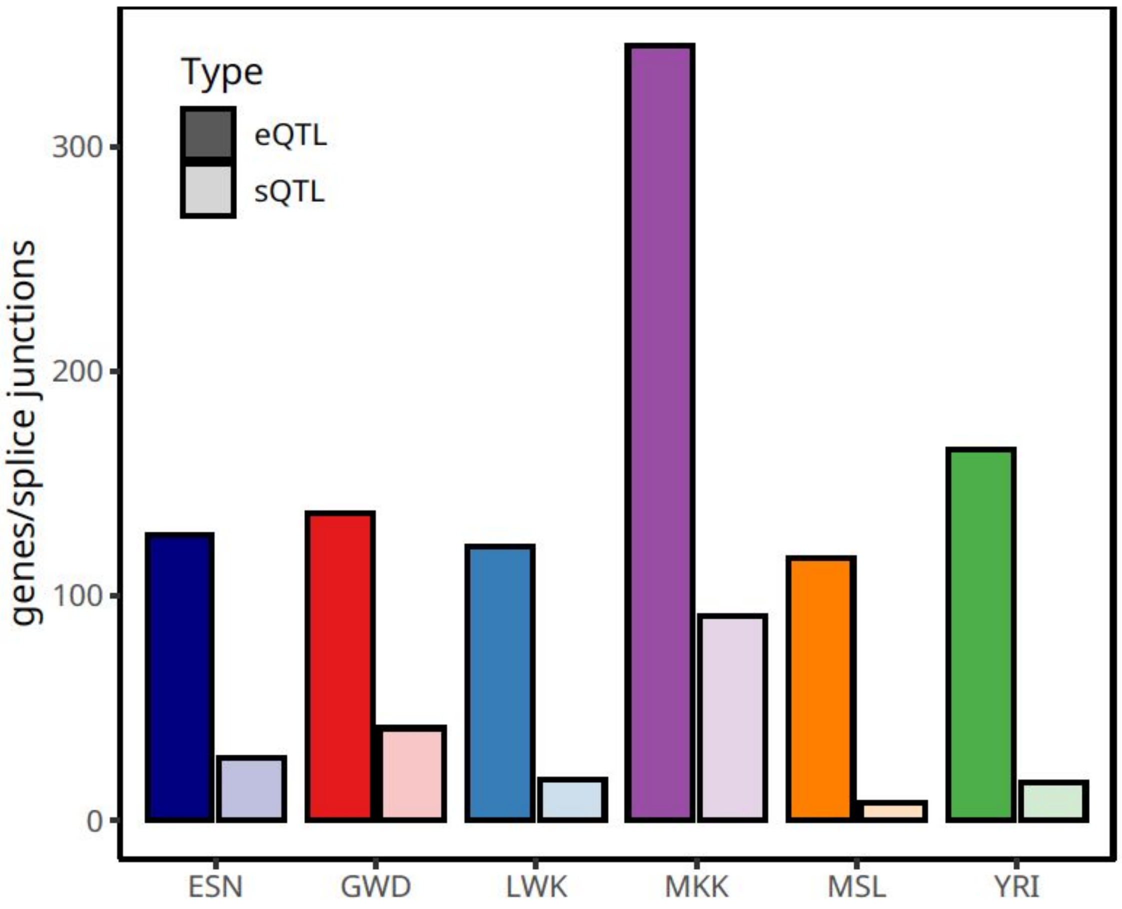


Supplementary Figure 1. Number of population-specific eQTL and sQTL effects compared to all other populations

Number of eQTL genes (dark shaded) or sQTL splice junctions (light saded) with mashr estimates of effect size significantly different in one population when compared with the other populations combined.

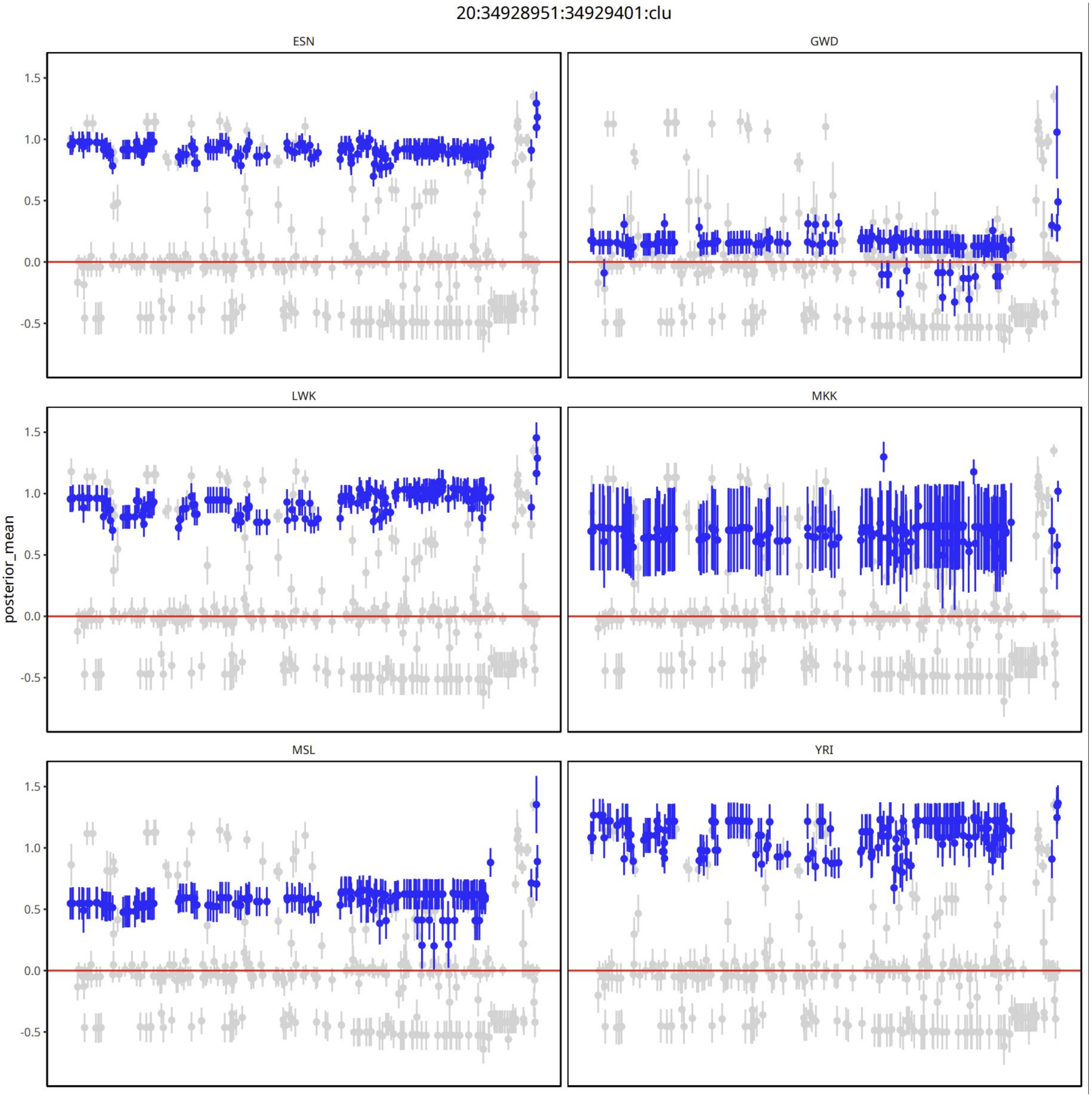


#### Supplementary Figure 2. Example of an sQTL signal in glutathione synthetase (GSS) with significantly different effect sizes across populations, as estimated by mashr

Variants are represented by their mashr-derived posterior z-scores (y-axis) and genomic position on chromosome 20 (x-axis). Variants with significantly different effect sizes between at least two populations are highlighted in blue. Effect sizes that were not different between populations are in grey.

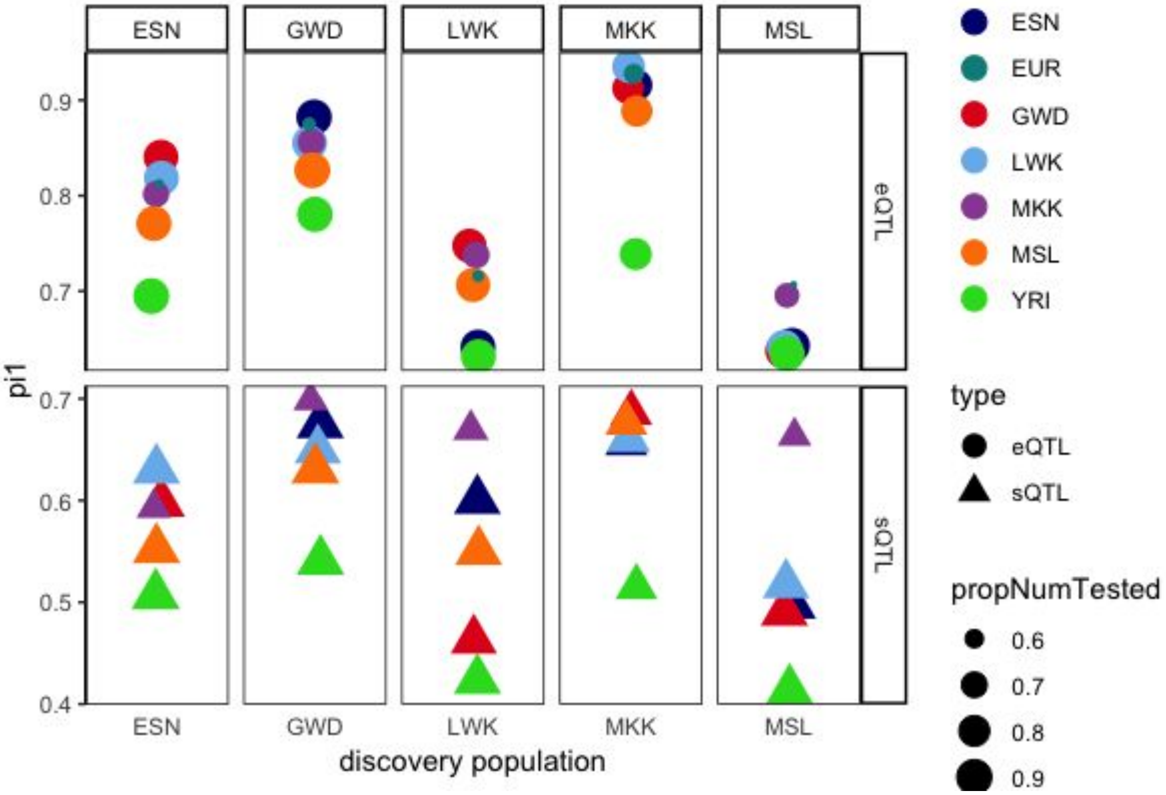


#### Supplementary Figure 3. eQTL and sQTL pi1

Pi1 statistic of replication of top 3000 eQTL (top) and top 3000 sQTL (bottom) from each population in the other AFGR populations, and in European populations from the GEUVADIS project. The size of the point represents the proportion of top variants that were tested in both populations.

a

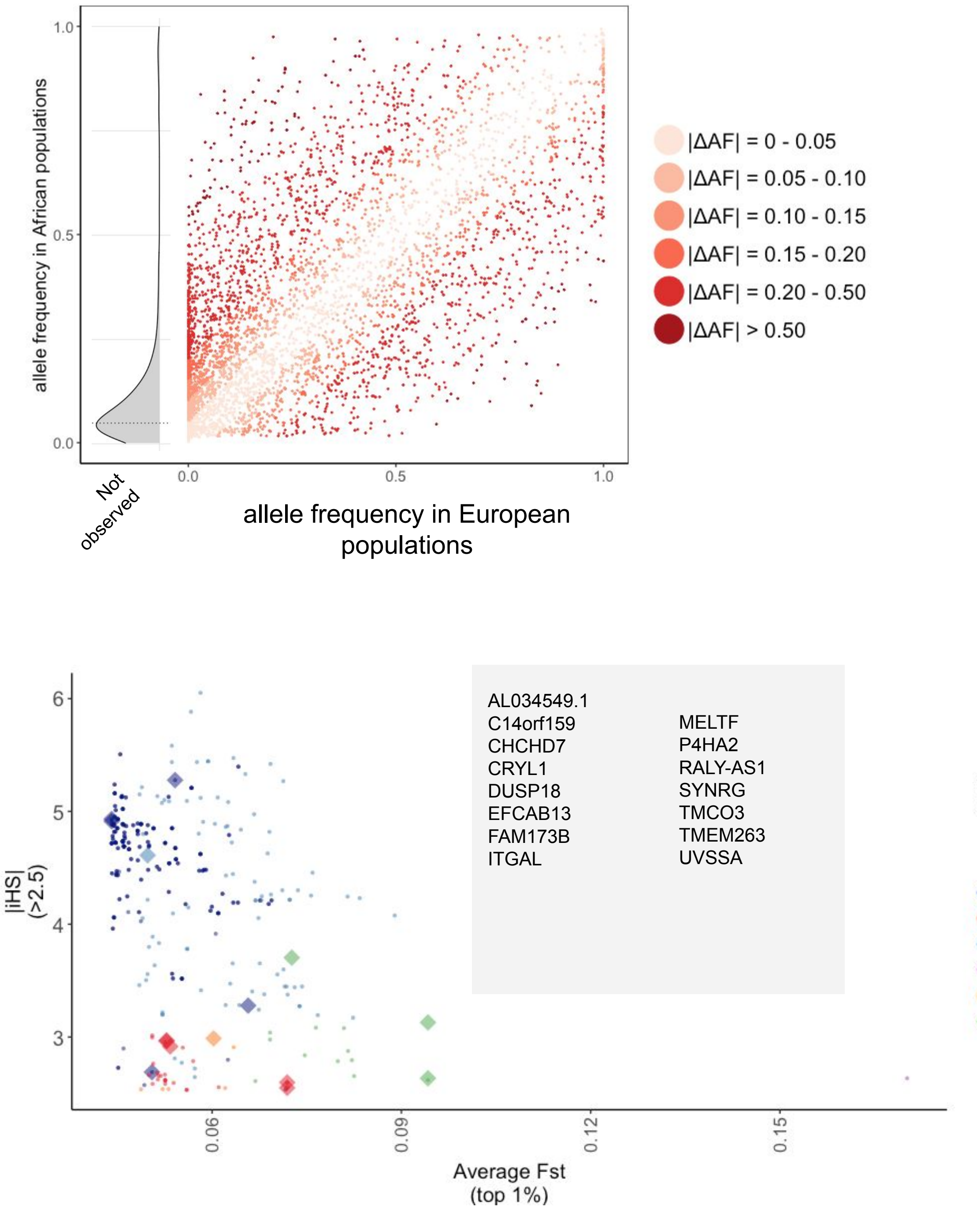


Supplementary Figure 4. Continental allele frequency differences and evidence of selection at sQTL variants

a. Allele frequency distribution of lead AFGR meta-analysis sQTL variants in European populations from the 1000 Genomes Project. The allele frequency distribution in AFGR of lead meta-analysis variants not present in European populations (2158) is depicted in gray at the far left of the plot.

b. Lead variants (diamond), and variants in high ld (r2 > 0.8) with lead variants (circle), from population-level sQTL with |iHS| > 2.5 and top 1% average Fst scores.

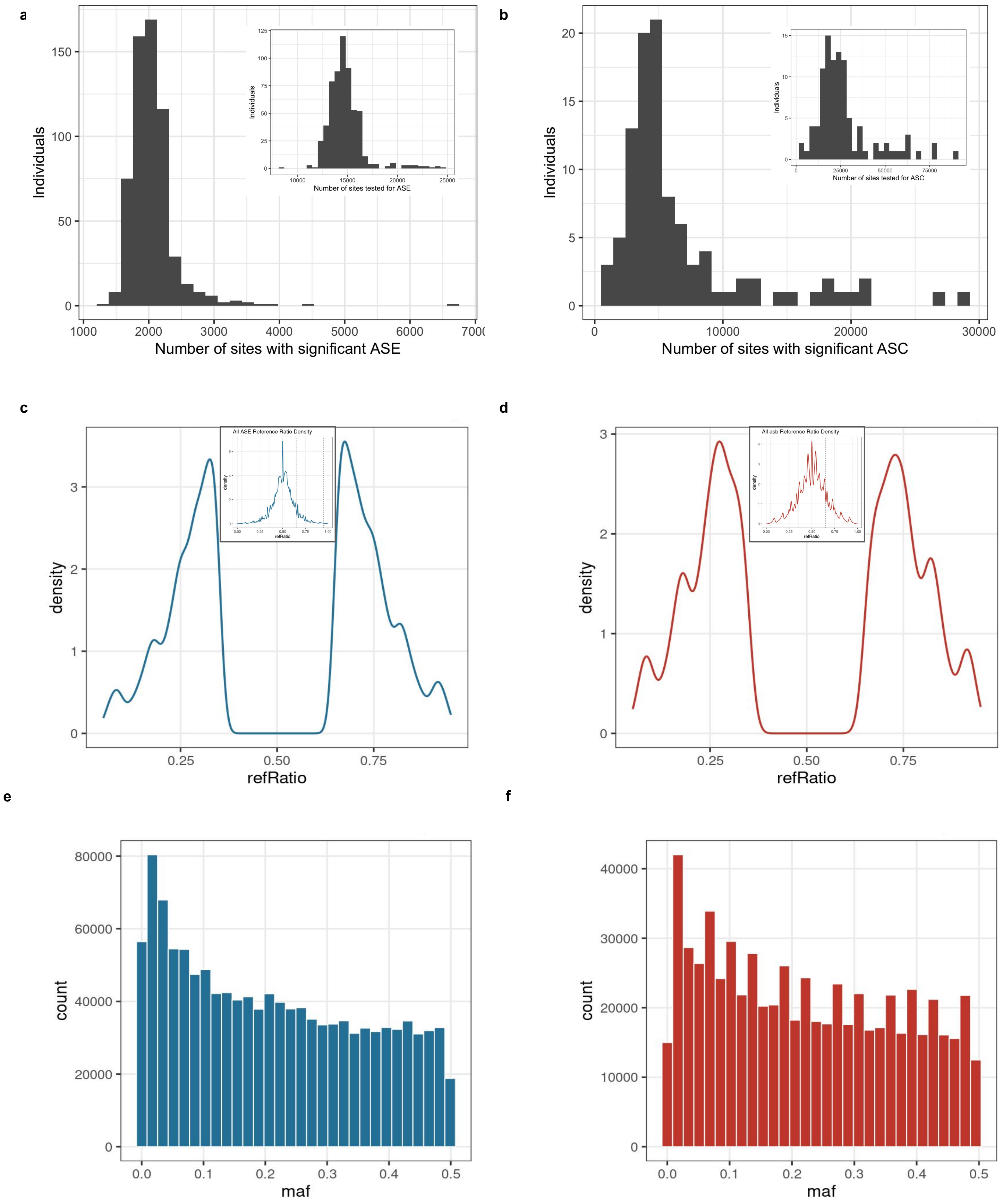
a
b
c
d
e
f
Individuals
Number of sites with significant ASE
Number of sites tested for ASE
Number of sites with significant ASC
Number of sites tested for ASC
All ASE Reference Ratio Density
All asb Reference Ratio Density
density
refRatio
count
maf

#### Supplementary Figure 5. ASE and ASB summary stats

a. Number of significant ASE variants detected per sample; inset displays the number of variants tested for ASE.
b. Number of significant ASC variants detected per sample; inset displays the number of variants tested for ASC.
c. Density plot of the reference ratios at significant ASE sites (main plot) and at all tested ASE sites (inset)
d. Density plot of the reference ratios at significant ASC sites (main plot) and at all tested ASC sites (inset)
e. Minor allele frequency spectrum for significant ASE sites
f. Minor allele frequency spectrum for significant ASC sites

### S6. RNA-seq quality control: expression

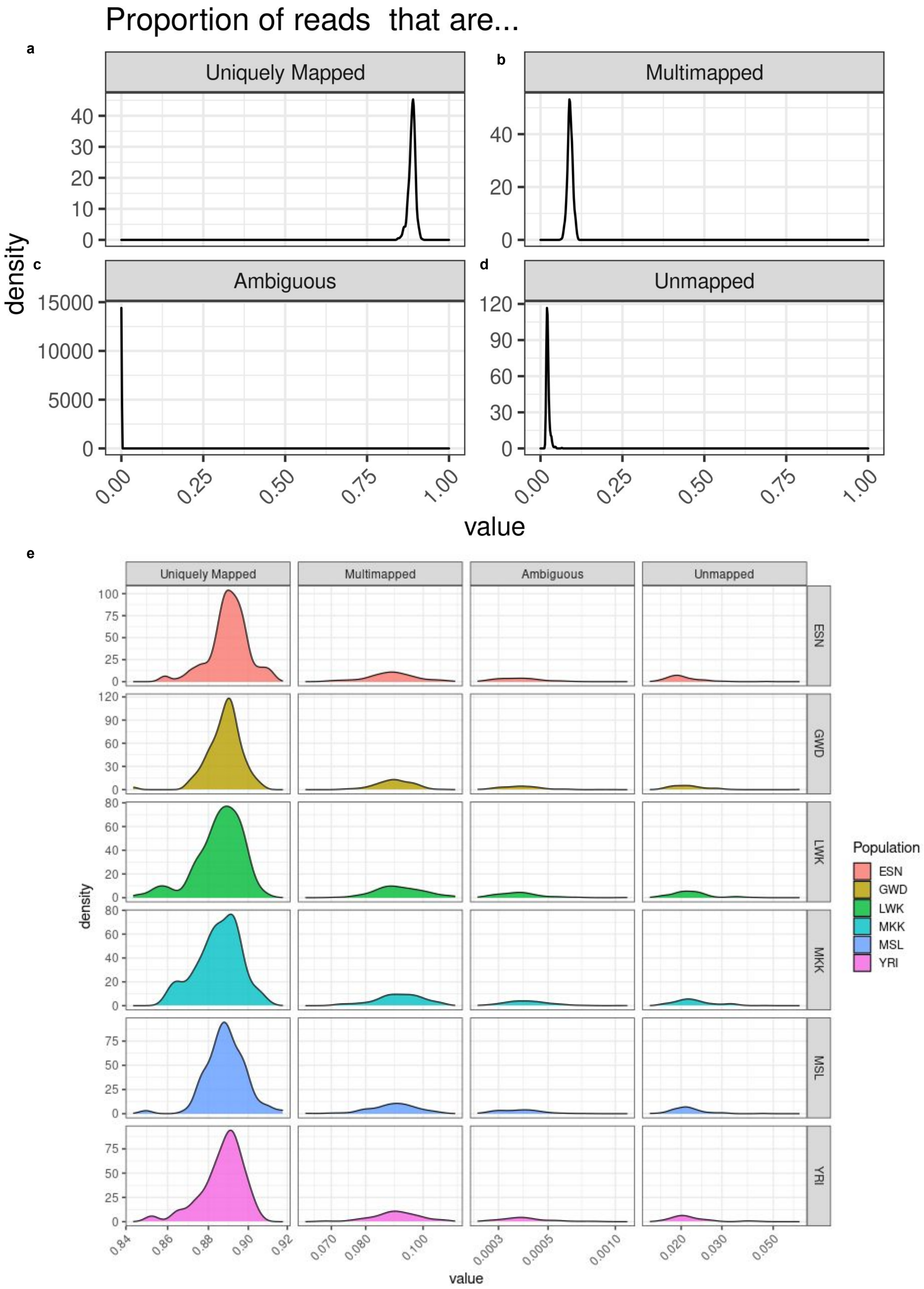

#### Supplementary Figure 6. RNA-Seq quality control: expression

RNA-seq alignment metrics from STAR aligner showing the proportion of total reads across all AFGR populations

a. mapped to a single genomic region
b. mapped to multiple genomic regions
c. had ambiguous mapping, or
d. that were unmapped.
e. proportion of reads by mapping status per population.

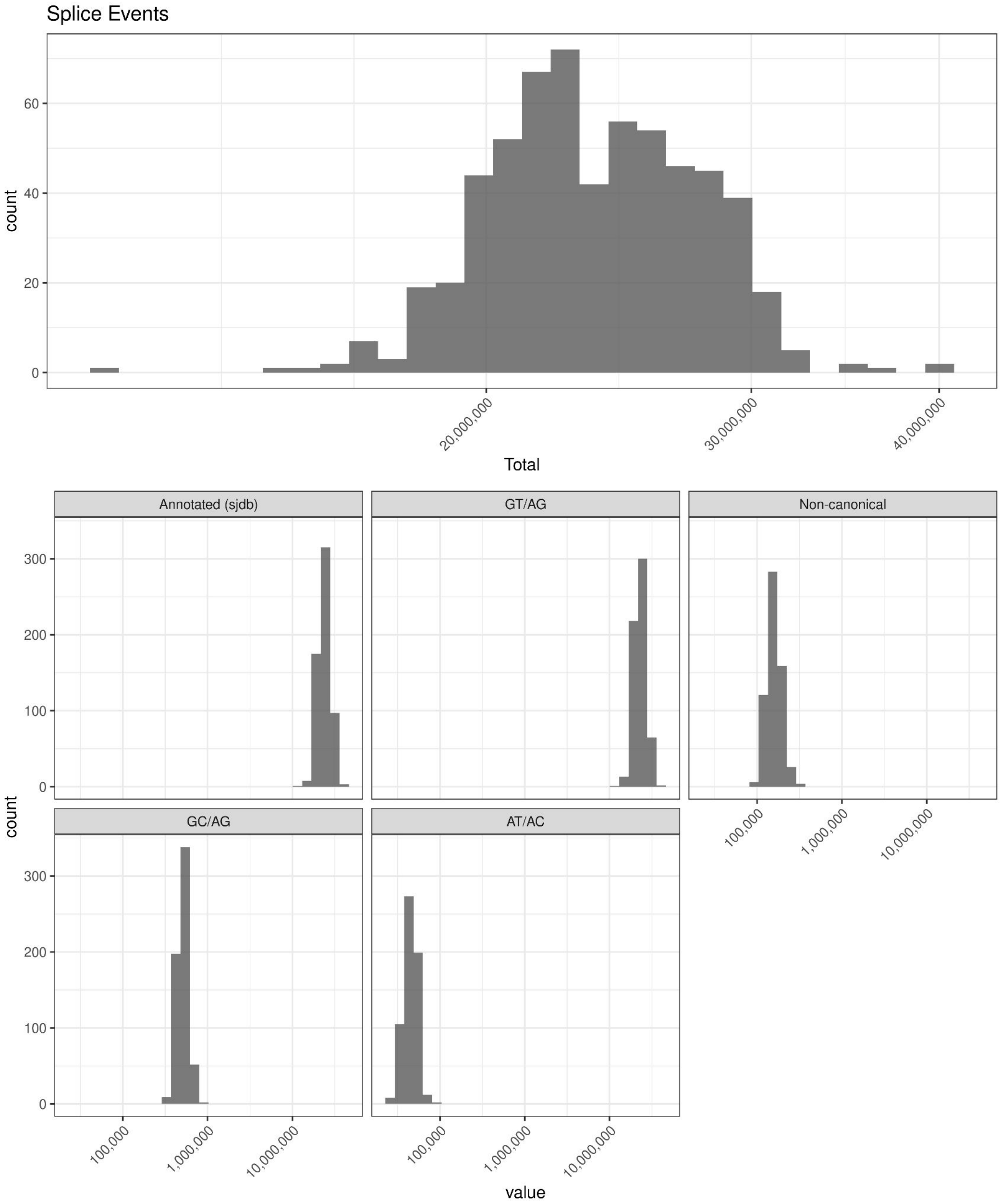


### Supplementary Figure 7. RNA-Seq quality control: splicing

a. Total number of splice events per sample detected across AFGR
b. Number of annotated, canonical, and non-canonical splice junctions per per sample across all AFGR samples

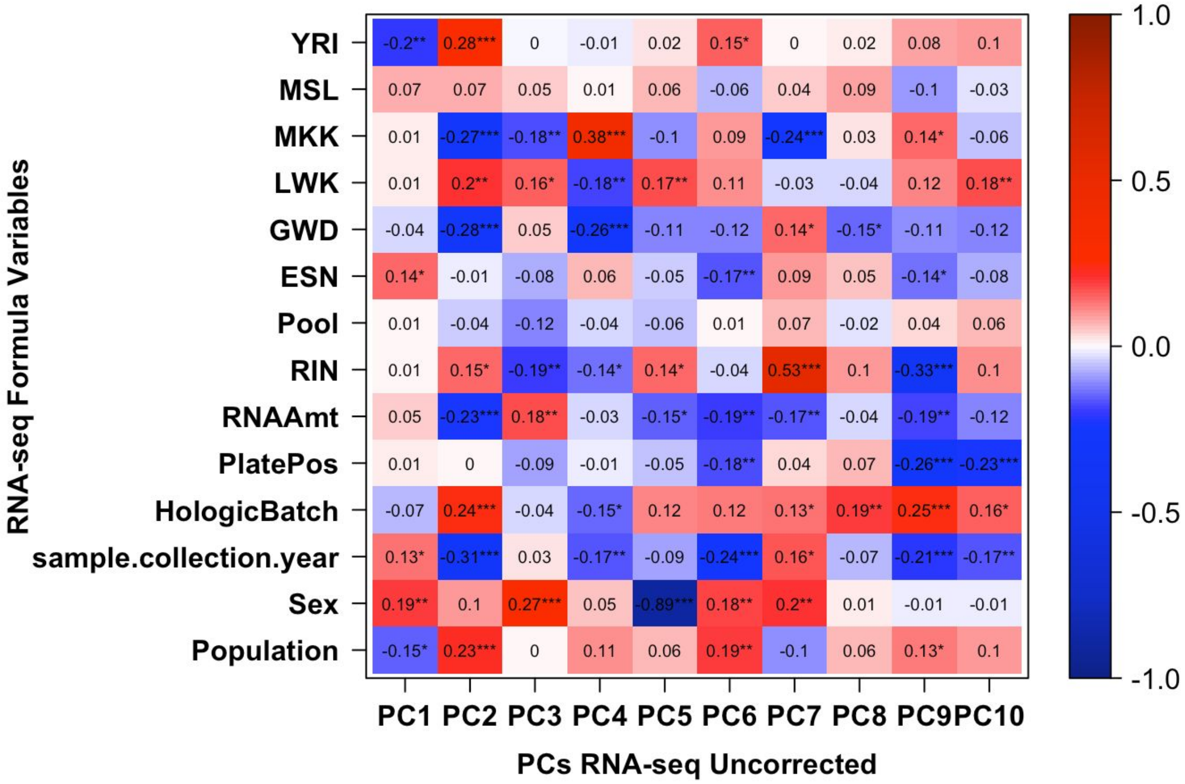


Supplementary Figure 8. RNA-Seq principal components associated with known variables

Heatmap depicting the correlations between the top 10 RNA-seq PCs and covariates. Scale and labels indicate the pearson correlation coefficient *r*, with asterix indicating the significance of the association: *** p-val < 0.001; ** p-val < 0.01; * p-val < 0.05.

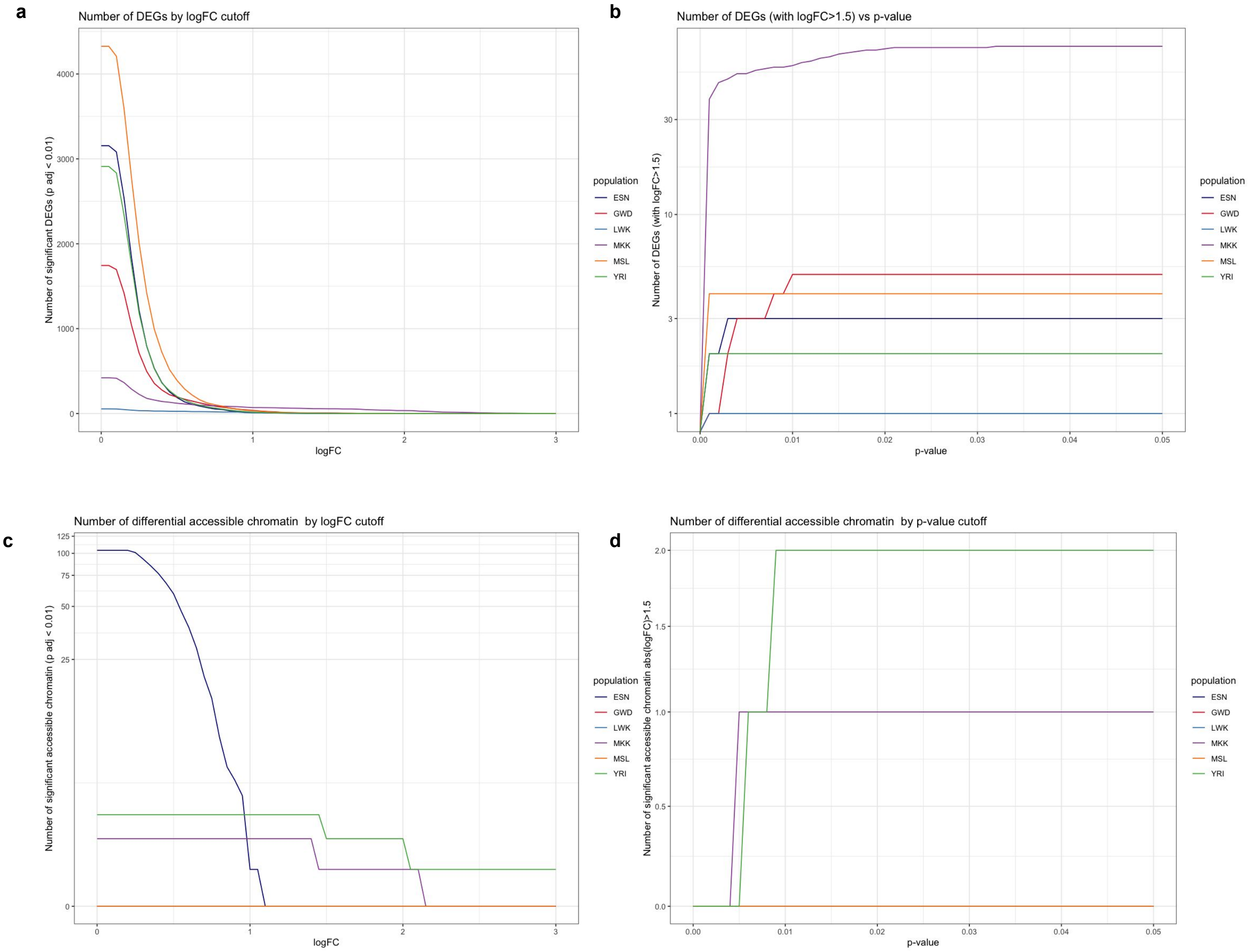


### Supplementary Figure 9. Differential gene expression logFC cutoffs

a. Number of significant differentially expressed genes identified per population at a constant adjusted p-value threshold of 0.01 and an absolute logFC cutoff ranging from 0 to 3.
b. Number of significantly differentially expressed genes detected per population at an absolute logFC cutoff of 1.5 and a maximum adjusted p-value threshold ranging from 0 to 0.5.
c. Number of significant differentially accessible chromatin regions identified per population at a constant adjusted p-value threshold of 0.01 and an absolute logFC cutoff ranging from 0 to 3.
d. Number of significantly differentially accessible chromatin regions detected per population at an absolute logFC cutoff of 1.5 and a maximum adjusted p-value threshold ranging from 0 to 0.5.

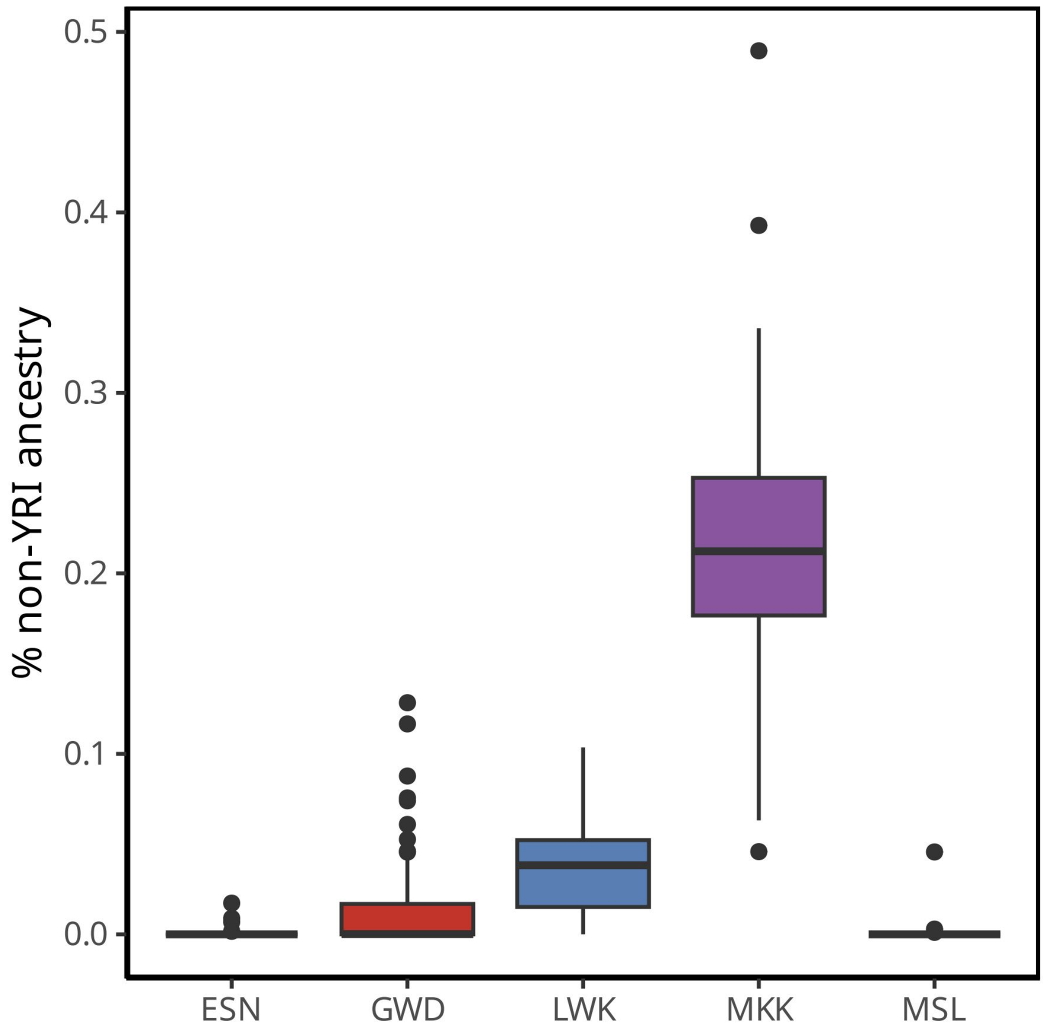


Supplementary Figure 10. Global ancestry: percent non-West African (YRI) ancestry detected genome-wide per population

Percent of non-West African ancestry detected genome-wide, using West African (YRI), European (CEU, TSI), East Asian (CHB, JPT), and South Asian (BEB, STU) reference populations from the 1000 Genomes Project.

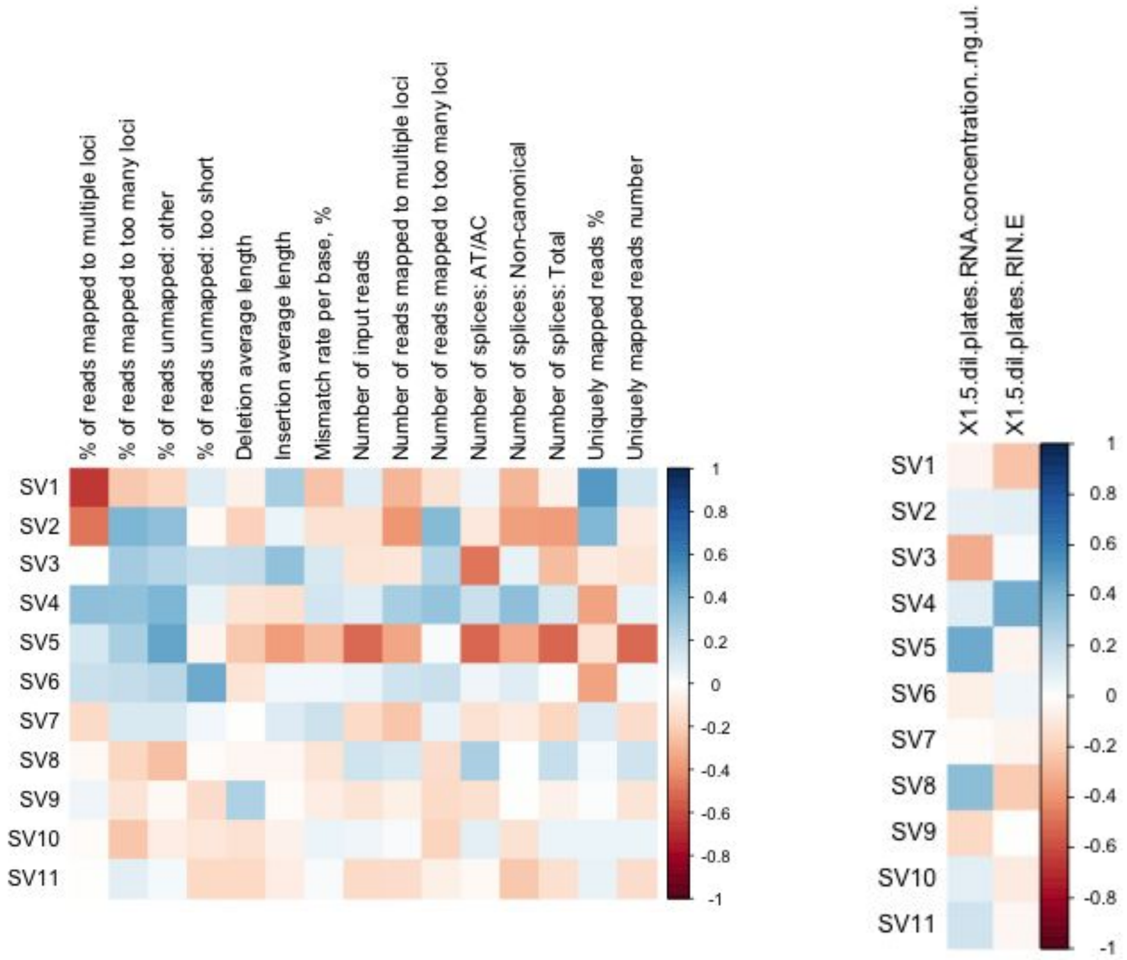


Supplementary Figure 11. RNA-Seq eQTL surrogate variables correlations with known variables - ESN

Correlation of surrogate variables included when calling eQTLs in ESN with other known variables. Surrogate variables were calculated using SVA.

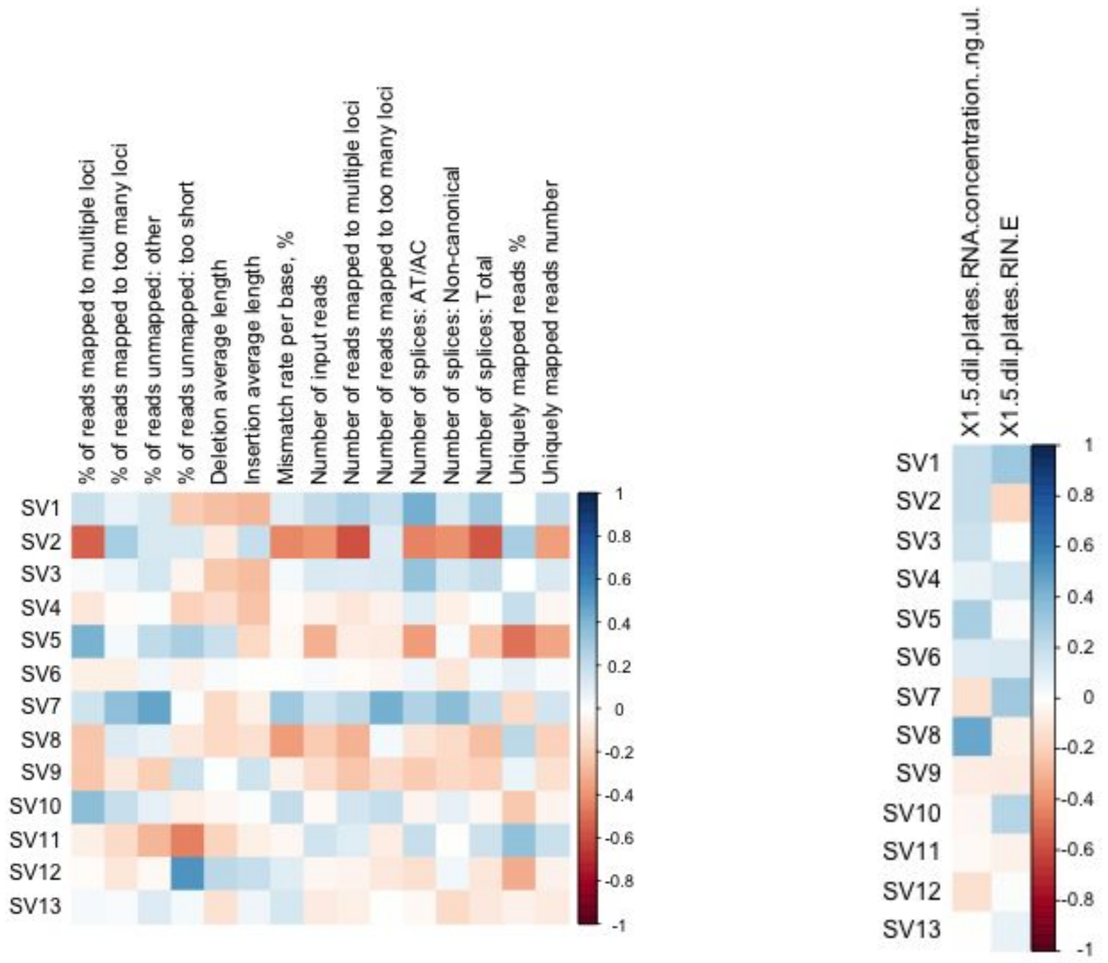


#### Supplementary Figure 12. RNA-Seq eQTL surrogate variables correlations with known variables - GWD

Correlation of surrogate variables included when calling eQTLs in GWD with other known variables. Surrogate variables were calculated using SVA.

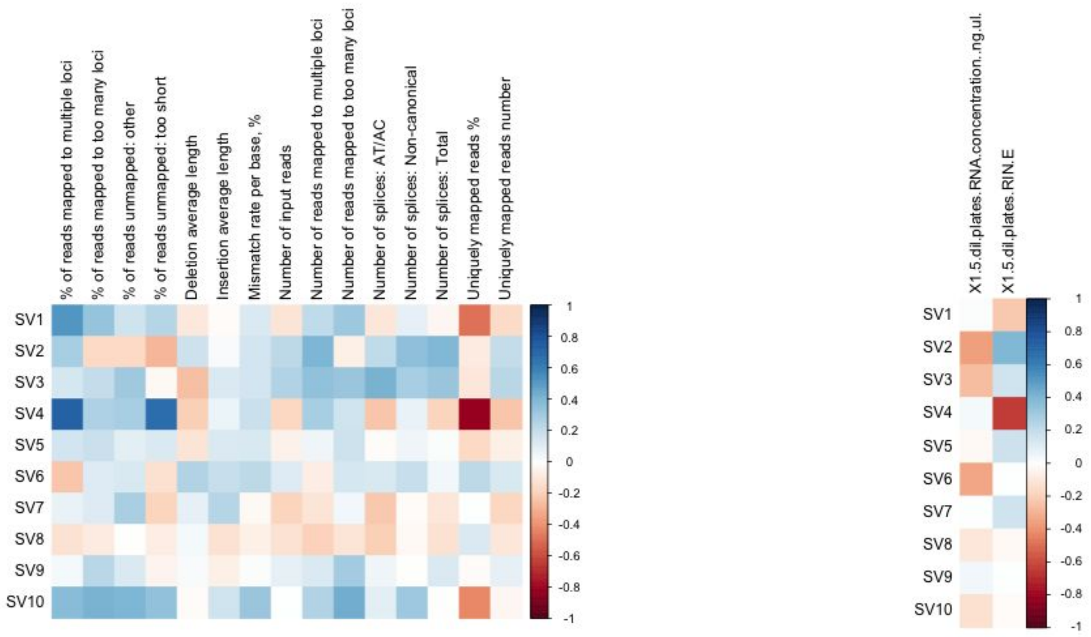


#### Supplementary Figure 13. RNA-Seq eQTL surrogate variables correlations with known variables - LWK

Correlation of surrogate variables included when calling eQTLs in LWK with other known variables. Surrogate variables were calculated using SVA.

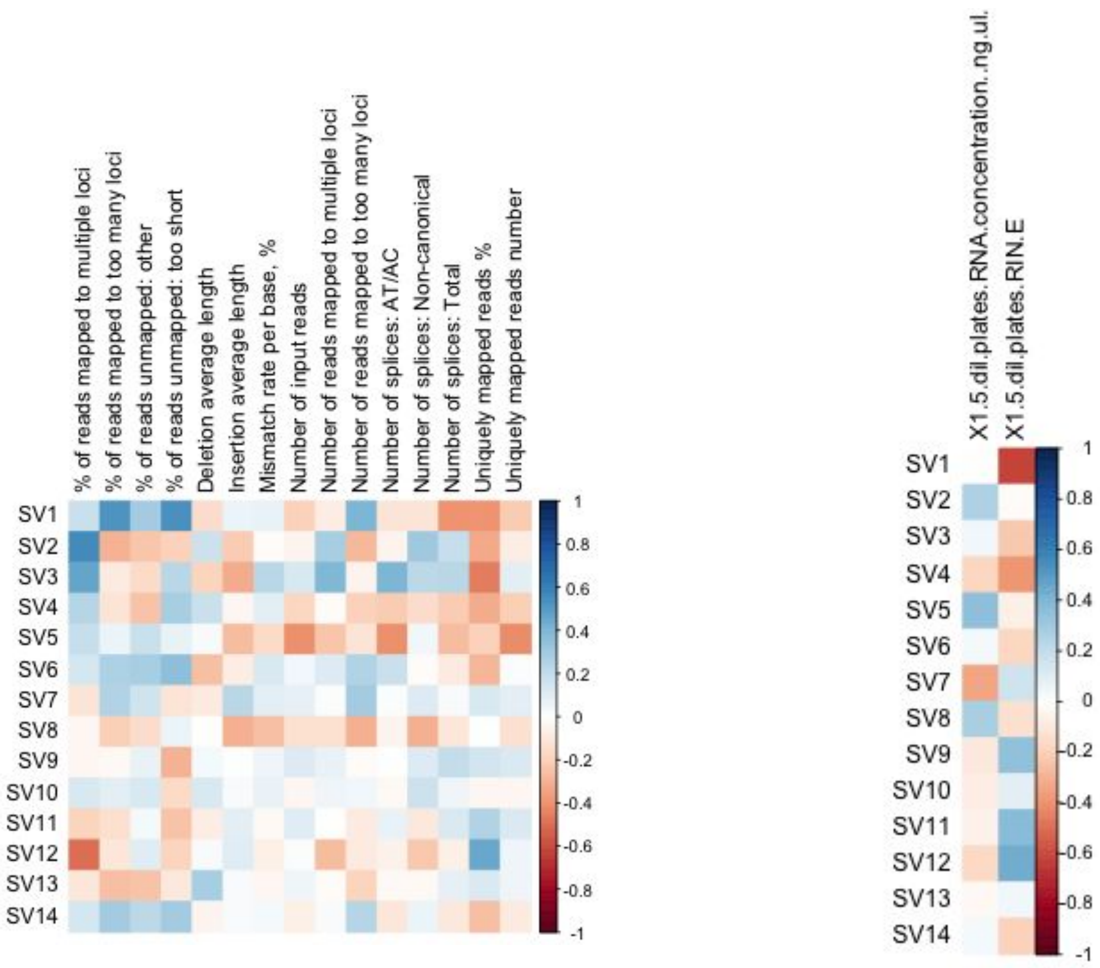


Supplementary Figure 14. RNA-Seq eQTL surrogate variables correlations with known variables - MKK

Correlation of surrogate variables included when calling eQTLs in MKK with other known variables. Surrogate variables were calculated using SVA.

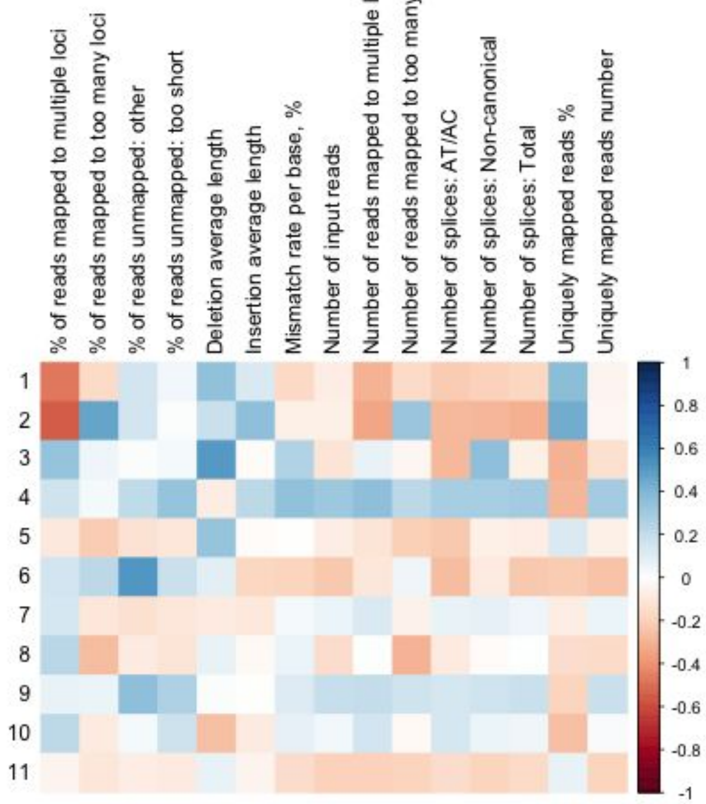


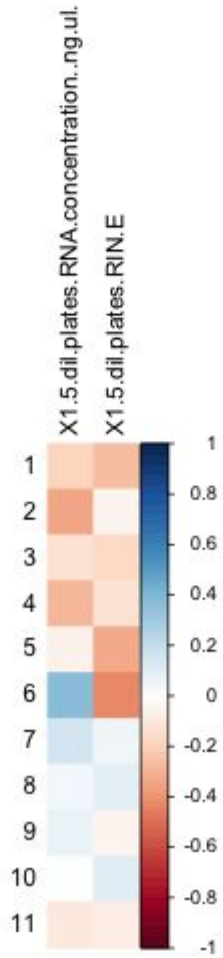


#### Supplementary Figure 15. RNA-Seq eQTL surrogate variables correlations with known variables - MSL

Correlation of surrogate variables included when calling eQTLs in MSL with other known variables.
Surrogate variables were calculated using SVA.

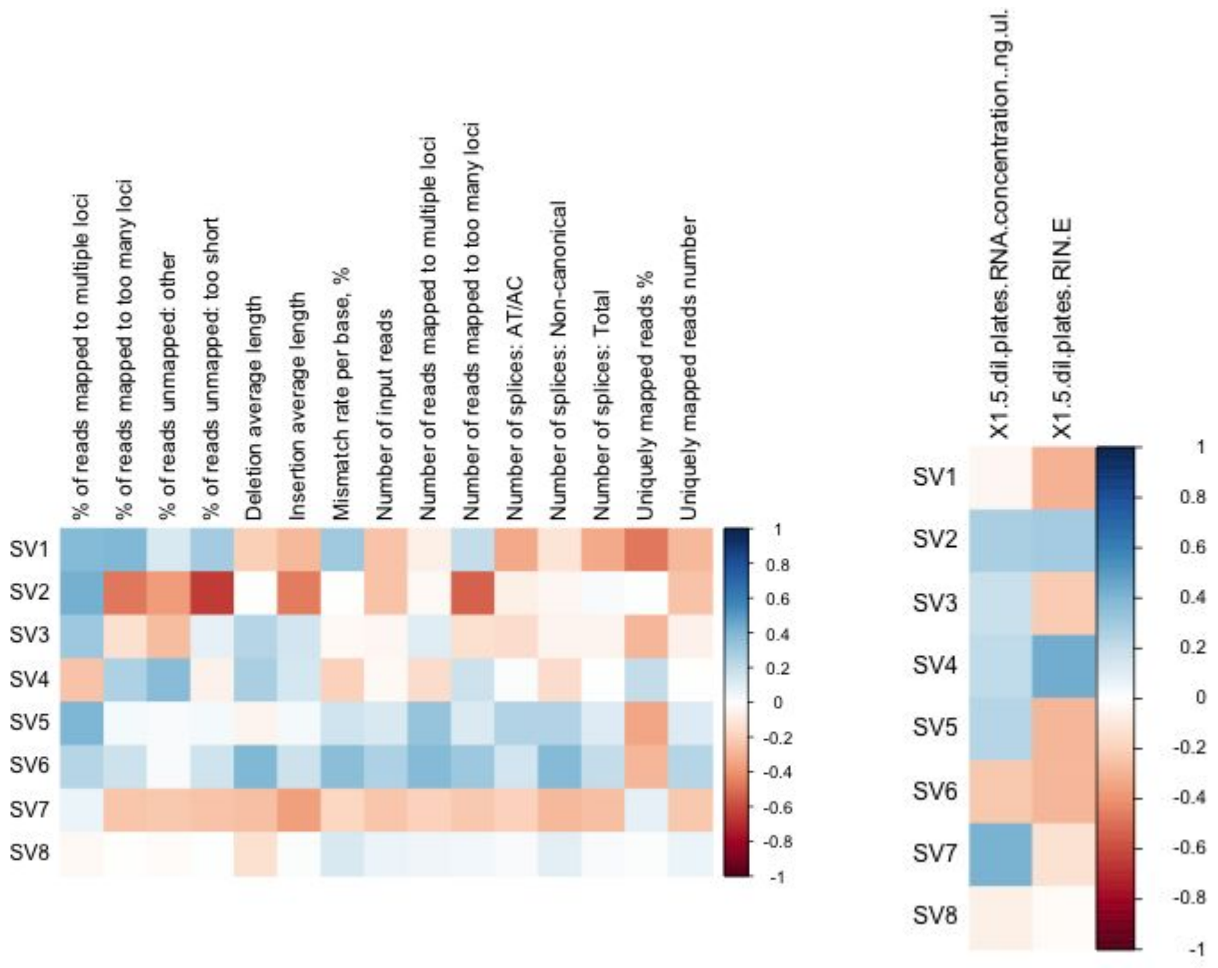


Supplementary Figure 16. RNA-Seq eQTL surrogate variables correlations with known variables - YRI

Correlation of surrogate variables included when calling eQTLs in YRI with other known variables. Surrogate variables were calculated using SVA.

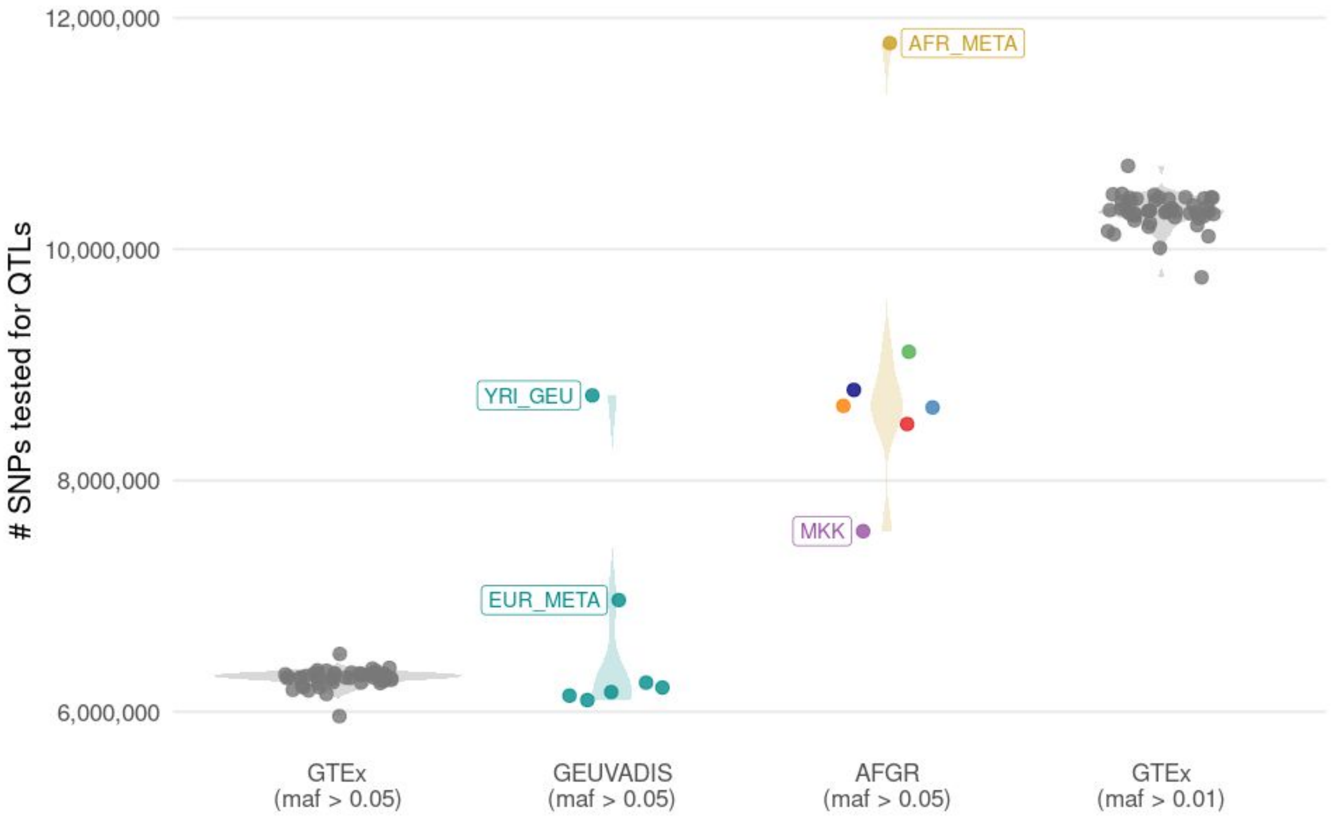


#### Supplementary Figure 17. number of variants tested for eQTLs in multiple large cohorts

Number of variants with allele frequencies (maf > 0.01 or maf > 0.05) testable for (e)QTL associations in functional genomics datasets from GTEx (Genotype-Tissue Expression project), GEUVADIS (Genetic European Variation in Disease consortium), and AFGR (African Functional Genomics Resource).

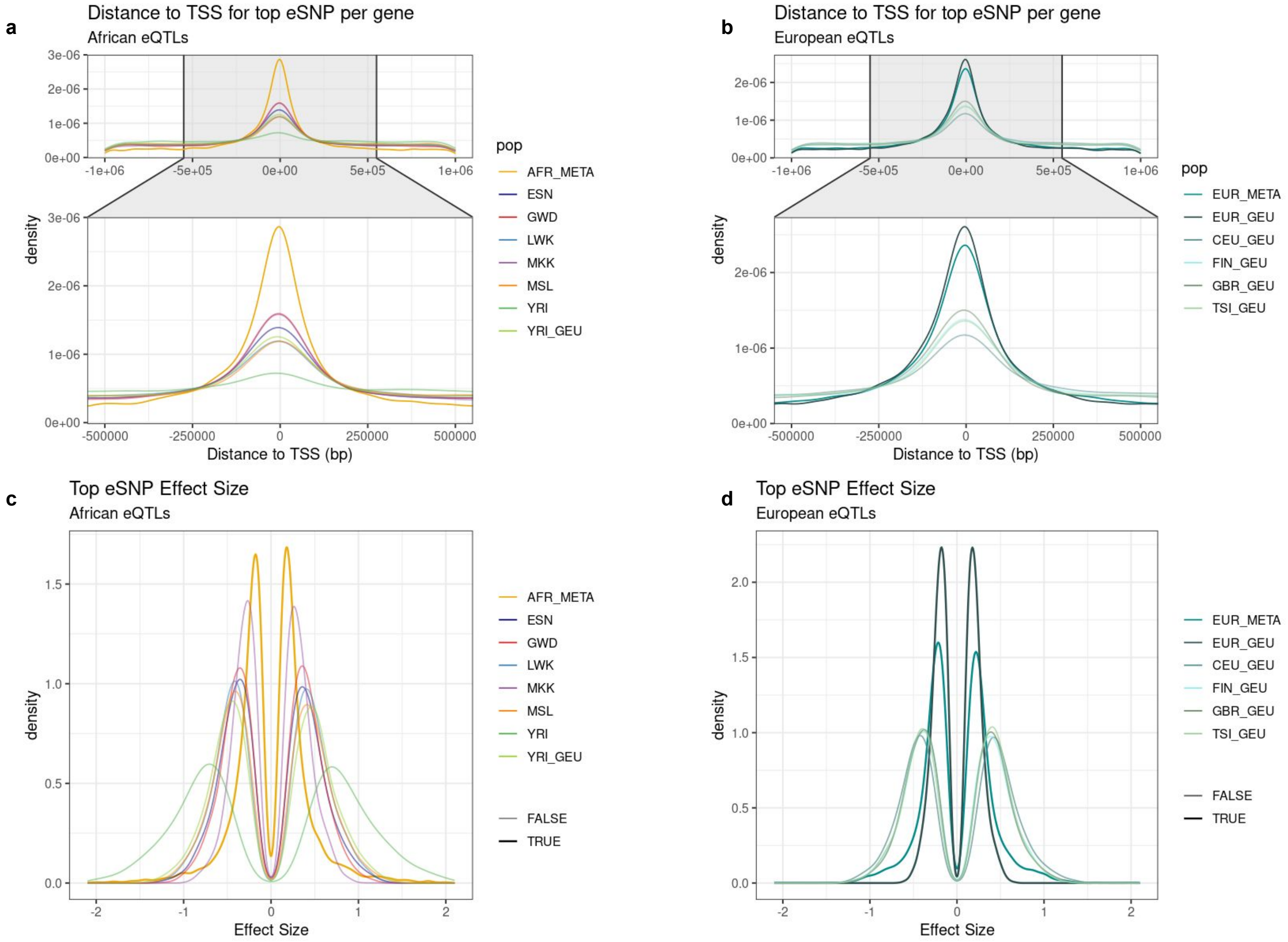


S18. eQTL Meta-analysis quality control

a. Distance from the top eSNP per gene to the TSS in the African meta-analysis and AFGR population eQTLs
b. Distance from the top eSNP per gene to the TSS in the European meta-analysis and GEUVADIS European population eQTLs
c. Effect size distribution for the top eSNP per gene in the African meta-analysis and African populations
d. Effect size distribution for the top eSNP per gene in the European meta-analysis and European populations

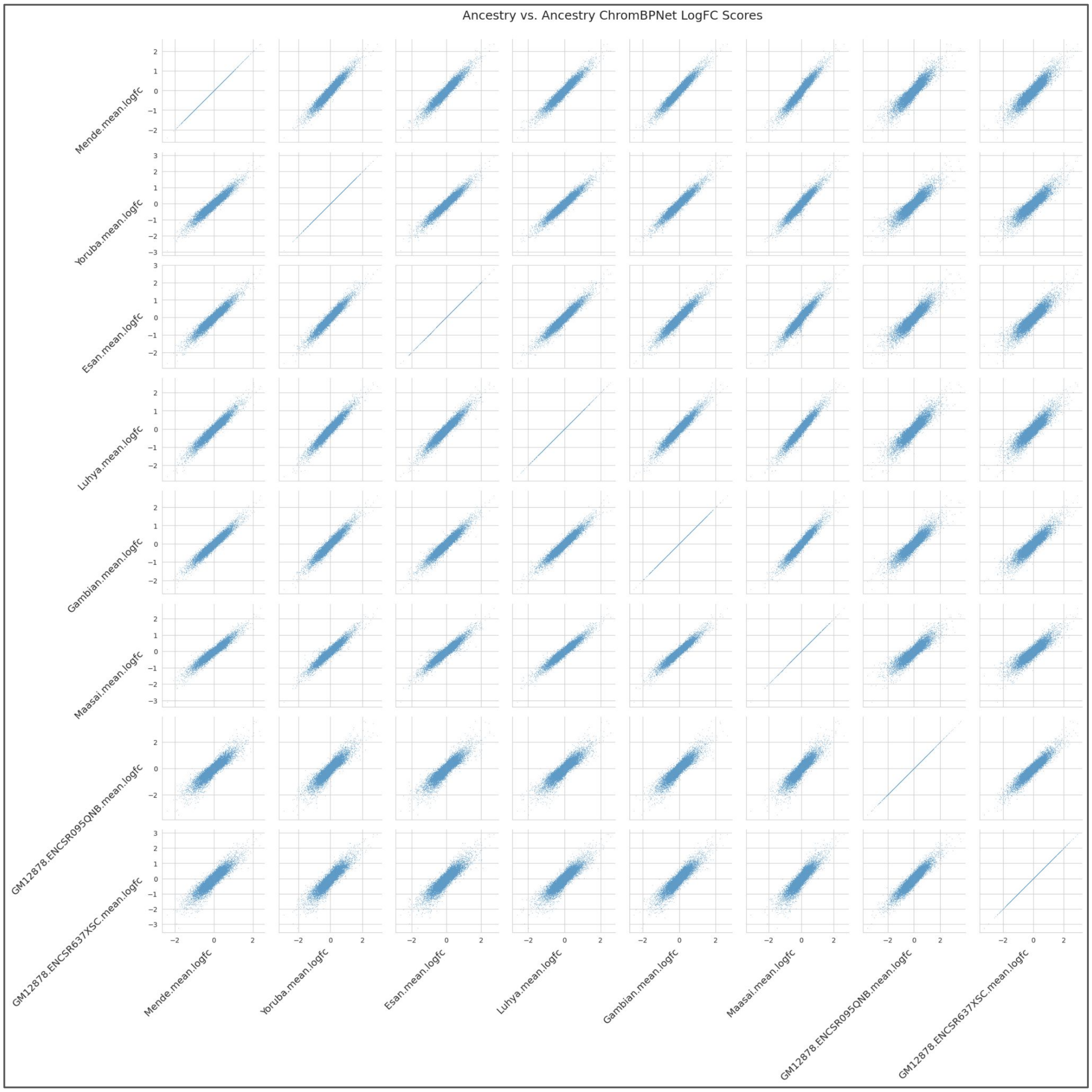


#### Supplementary Figure 19. Pairwise correlation of chromBPnet logFC prediction models

Mean logFC scores for each African population-derived model (ESN, GWD, LWK, MKK, MSL, YRI) and two GM12878-derived models, correlated pair-wise with mean logFC predictions from all other models, for all caQTL SNPs in peaks (n=219,382).
