## Supplementary Tables for "Transcriptomics and chromatin accessibility in multiple African population samples": biorxiv-AFGR-SupplementaryTableS2.pdf

Supplementary Table 2. Number of transcribed features detected across all AFGR samples. Unnanotated features do not have an annotated in GENCODEv27, and novel features are a subset of the unannotated that are also not detected in long-read RNA-seq of GM12787 or GENCODEv43. African pan-genomic contig features are those for which reads preferentially aligned to the a contig from the pan-genomic contig collection published by Sherman et al.

| **Feature** | **Reference genome expression** | | | **African pan-genomic contig expression** |
| --- | --- | --- | --- | --- |
| | Total | Unannotated | Novel | Novel |
| Exons | 310504 | 57916 | 43994 | 2072 |
| Transcripts | 108971 | 26521 | 25277 | 367 |
| Loci | 21021 | 1303 | 248 | 284 |
