## Supplementary Tables for "Transcriptomics and chromatin accessibility in multiple African population samples": biorxiv-AFGR-SupplementaryTableS10.pdf

Supplementary Table 10. Splice Cluster Filtering

| Step | Clusters | Introns |
| --- | --- | --- |
| Leaf Cutter output (clustered across all African populations) | 37,490 (min 2, median 3, max 72 introns/cluster) | 151,808 |
| Filter out clusters with more than 10 introns | 35,798 | 124,844 |
| By population, introns supported by at least 10% of total number reads assigned to the cluster in at least 25% of samples, then intersect to set that passes in every population | 32,609 | 56,920 |
| Filter out clusters with fewer than 2 active introns (14,812 single intron clusters) | 17,797 | 42,108 |
| Filter out clusters with low splicing variability, ie, Hellinger's distance < 0.01 | 13,359 | 32,493 |
